# *Seda*DNA metabarcoding reveals hidden histories of within-species sequence variation

**DOI:** 10.64898/2026.11.03.763919

**Authors:** Elena Baños, Jesús Zarcero, Andrew B. Cundy, Enrique González-Ortegón, Erik J. de Boer, Xavier Benito, Xavier Turon, Luke E. Holman, Sandra Nogué, Marc Rius

**Affiliations:** Department of Marine Ecology, Centre for Advanced Studies of Blanes (CEAB), Spanish Research Council (CSIC), 17300 Blanes, Catalonia, Spain; Facultat de Biologia, Universitat de Barcelona, 08028 Barcelona, Catalonia, Spain; Departament de Biologia Vegetal, Biologia Animal i d’Ecologia (BABVE), Universitat Autònoma de Barcelona, Bellaterra (Cerdanyola del Vallès), 08193, Catalonia, Spain; Geological Institute, Department of Earth Sciences, ETHZ, Zurich, Switzerland; GAU-Radioanalytical, School of Ocean and Earth Science, National Oceanography Centre Southampton, University of Southampton, Southampton, United Kingdom; Stable Isotope Laboratory, School of Ocean and Earth Science, National Oceanography Centre Southampton, University of Southampton, Southampton, United Kingdom; Department of Ecology and Coastal Management, Instituto de Ciencias Marinas de Andalucía (ICMAN-CSIC), Spanish National Research Council (CSIC), 11510 Puerto Real, Cádiz, Spain; CREAF, Bellaterra (Cerdanyola del Vallès), 08193, Catalonia, Spain; IRTA, La Ràpita (Tarragona), 43540, Spain; Section of Biological Sciences, Institute of Catalan Studies (IEC), 08001 Barcelona, Catalonia, Spain; Section for Molecular Ecology and Evolution, Globe Institute, University of Copenhagen, Copenhagen, Denmark; Department of Zoology, Centre for Ecological Genomics and Wildlife Conservation, University of Johannesburg, Auckland Park Johannesburg 2006, South Africa

## Abstract

Most genetic data come from contemporary populations, leaving historical changes in within-species variation poorly documented. Sedimentary ancient DNA (*seda*DNA) offers an opportunity to recover historical sequence variation, but its potential to reconstruct temporal changes in within-species composition remains underexplored. Here, we analysed dated sediment cores from eight Atlantic and Mediterranean coastal wetlands using COI and 18S metabarcoding, treating amplicon sequence variants as putative haplotypes. Across native, non-indigenous and cryptogenic taxa, accumulated haplotype diversity and geographic sharing varied markedly among species and markers, without consistently distinguishing species status. Temporal reconstructions revealed that accumulated haplotype networks combined distinct sets of variants detected across historical periods. Broader analyses of 18S data further showed substantial variation in haplotype composition among periods, sites and taxa. By resolving the temporal and spatial distributions of within-species sequence variants, our findings demonstrate how sedimentary DNA archives can uncover historical dimensions of biodiversity obscured by time-aggregated analyses, establishing a foundation for historical metaphylogeography.

## Introduction

Genetic variation within species changes through time as populations respond to demographic processes, dispersal and environmental change (Habel et al., 2014). Yet much of what we know about this variation comes from contemporary samples, which provide only a limited view of its history (Clark et al., 2025; Wandeler et al., 2007). Present-day genetic patterns may conceal earlier differences in the identity and geographic distribution of variants, making it difficult to establish how within-species diversity has changed (Ramakrishnan & Hadly, 2009). Historical genetic records are therefore important not only for documenting past diversity, but also for understanding how contemporary patterns have emerged.

Sedimentary ancient DNA (*seda*DNA) offers a means of extending biodiversity observations into the past. Dated sedimentary archives have provided records of plants, animals and microorganisms across terrestrial, freshwater and marine ecosystems (Crump, 2021; Kjær et al., 2022; O’Reilly-Berkeley et al., 2026; Zimmermann et al., 2023). These records have revealed changes in vegetation and ecological communities associated with climatic variation and human activities (Alsos et al., 2024; Ficetola et al., 2018; Garcés-Pastor et al., 2022). Most sedimentary DNA metabarcoding studies, however, have focused on species occurrences or community composition. Less attention has been given to how sequence variation within species is distributed across the periods represented by these archives.

Metabarcoding can recover distinct amplicon sequence variants (ASVs) assigned to the same species, which can be examined as putative haplotypes when marker resolution and taxonomic assignments are sufficiently reliable (Serrana & Watanabe, 2023; Tsuji et al., 2019). Previous sedimentary DNA studies have recovered historical within-species sequence variation and documented changes in detected variants through time (Epp et al., 2010; Kyalo-Omamo et al., 2023; Lammers et al., 2024). However, less is known about how within-species sequence variants recovered from sedimentary archives are distributed across historical periods and geographic locations, particularly when multiple taxa are considered. Pooling detections into accumulated haplotype networks obscures this temporal information, while comparisons based on haplotype richness alone may overlook differences in sequence composition.

Introduced species provide a useful context for examining historical sequence variation, as founder events, repeated introductions and admixture can produce contrasting patterns of genetic diversity (Dlugosch & Parker, 2008; Rius & Darling, 2014). Comparing taxa with different species status classifications allows us to examine variation in historical sequence composition without assuming that species status reflects a particular demographic history.

Here, we examine within-species sequence variation recovered from dated sediment cores at eight Atlantic and Mediterranean coastal wetlands using COI and 18S metabarcoding. We compare accumulated putative haplotype diversity and geographic sharing among native, non-indigenous and cryptogenic taxa, reconstruct the variants detected in successive historical intervals, and quantify temporal changes in sequence composition. We ask whether time-aggregated networks obscure differences in the identity and geographic representation of variants, and what compositional analyses reveal beyond haplotype richness. Together, these analyses examine what temporal and geographic information adds to our understanding of within-species sequence variation recovered from sedimentary archives.

## Methods

### Study sites and sediment sampling

Sediment cores were collected from eight coastal wetlands along the Atlantic and Mediterranean coasts of the Iberian Peninsula (Fig. 1; Table S1). Three sites were located in the Santoña estuarine system in northern Spain: Cicero (CCO), Colindres (COL) and Santoña (STO). Four sites were located in southern Spain: two on Saltés Island, Huelva, along the Tinto River margin (HIT) and towards the open ocean (HIO), and two in the Bay of Cádiz, within saltmarshes in a restricted military area (CMB) and the Salinas Dolores complex (CSD). The remaining site was located in the Encanyissada lagoon of the Ebro Delta on the Mediterranean coast (DEE).

**Fig. 1:**
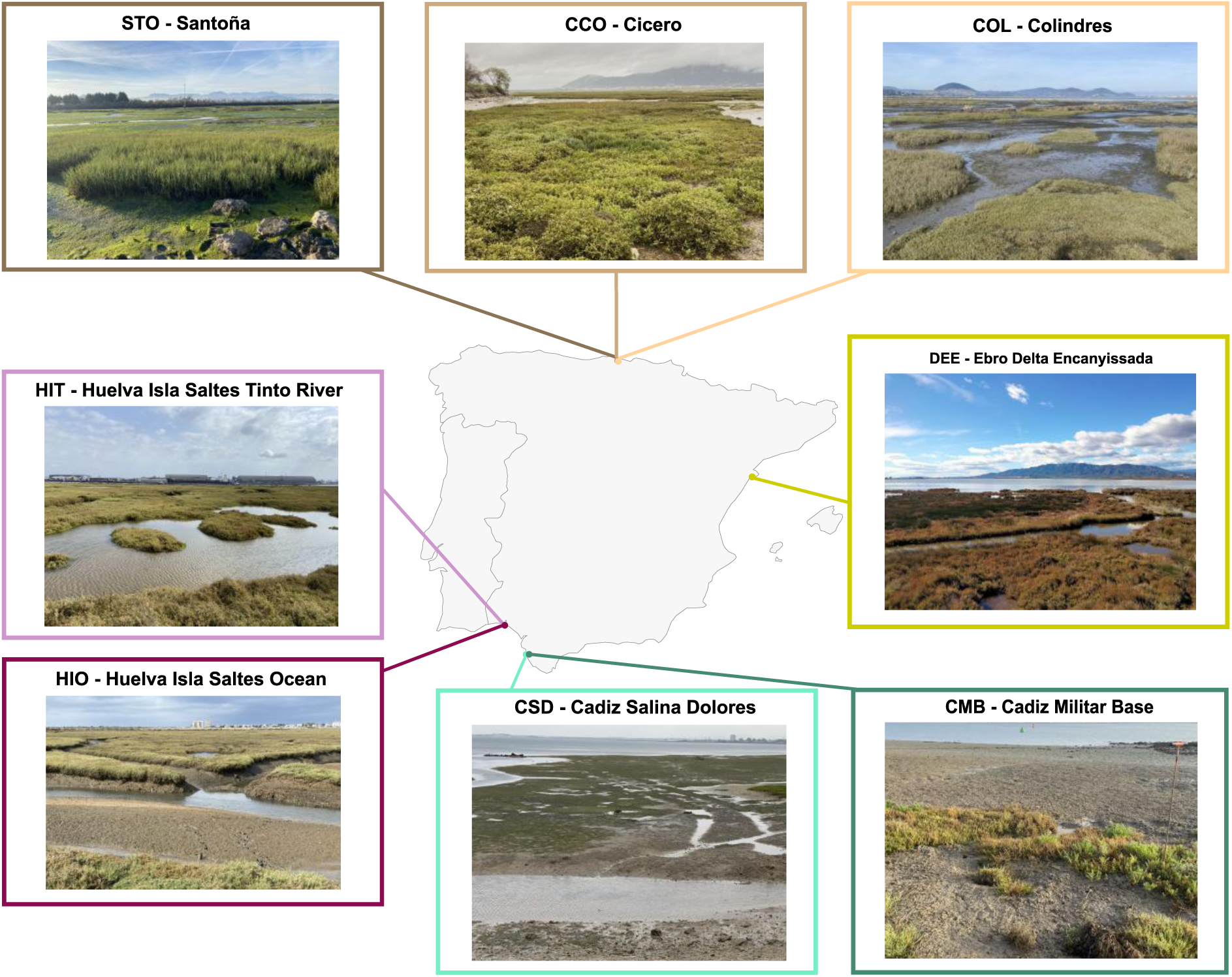
Location of the eight coastal wetland sampling sites across the Iberian Peninsula. The map shows the location of each site, accompanied by site photographs: Santoña (STO), Cicero (CCO), Colindres (COL), Cádiz Salina Dolores (CSD), Cádiz Militar Base (CMB), Huelva Isla Saltés–Tinto River (HIT), Huelva Isla Saltés–Ocean (HIO) and Ebro Delta Encanyissada (DEE).

Sediment cores were recovered using a Russian-type peat corer with a 50-cm sampling chamber, extending the sampling depth through successive insertions where necessary. Immediately after retrieval, cores were transferred to semicylindrical PVC tubes (5-cm internal diameter) previously decontaminated with bleach and rinsed with Milli-Q water. Cores were transported to the laboratory at 4 °C within 72 h.

Core opening and subsampling were conducted in a dedicated, positive-pressure, PCR-free laboratory physically separated from post-PCR facilities. Work surfaces and equipment were decontaminated with 10% sodium hypochlorite, rinsed with ultrapure water and treated with 70% ethanol. Before subsampling, approximately 3 mm of the outer sediment surface was removed to minimise contamination. Sediment was collected from the interior of the core, avoiding contact with the core walls. Subsamples of 2–5 g were collected from selected 1-cm depth intervals using sterile tools, with fresh sterile sampling materials used for each subsample. Sampling depths were spaced approximately 6 cm apart within the upper 60 cm and approximately 10 cm apart below this depth, with exact depths varying among cores. Subsamples were transferred to sterile 20-mL containers and stored at −20 °C until DNA extraction.

### Sediment chronology and temporal resolution

Sediment chronologies were investigated using lead-210 (²¹⁰Pb) dating at GAU-Radioanalytical, University of Southampton (UK), and accelerator mass spectrometry (AMS) radiocarbon (¹⁴C) dating at the Poznań Radiocarbon Laboratory (Poland). A total of 92 ²¹⁰Pb measurements were obtained across the eight sediment cores. Five additional samples from deeper sections of DEE, CSD, CMB, HIT and HIO were dated by AMS ¹⁴C to provide chronological information beyond the range of unsupported ²¹⁰Pb.

Recent sediment accumulation rates were estimated for each core using the Constant Flux– Constant Sedimentation (CF:CS) model (Krishnaswamy et al., 1971), based on linear regressions of ln-transformed unsupported ²¹⁰Pb activity against sediment depth. The lower limit of the ²¹⁰Pb-supported chronology was defined as the depth at which unsupported ²¹⁰Pb activity became indistinguishable from background activity. Calendar ages were estimated using the core-specific CF:CS accumulation rates. Ages below the unsupported-²¹⁰Pb dating horizons were obtained by linear extrapolation, assuming that the recent sediment accumulation rates remained constant over longer periods.

AMS ¹⁴C determinations were calibrated using the IntCal20 Northern Hemisphere atmospheric calibration curve (Reimer et al., 2020) with the R package *rcarbon* (v1.5.2; Crema & Bevan, 2021). No marine-reservoir or empirical old-carbon correction was applied. Because the dated carbon fraction was not independently characterised, calibrated radiocarbon ages were treated as estimates of the age of the dated carbon rather than direct estimates of sediment deposition. Accordingly, the radiocarbon determinations were retained as independent chronological comparisons and were not incorporated as depositional constraints into the working age–depth models.

To establish a common temporal framework across cores with different sediment accumulation rates and sampling resolutions, samples were assigned to 50-year intervals according to their estimated calendar ages (e.g., 1900–1949). These intervals were used throughout the temporal analyses. Chronological uncertainty was not explicitly propagated into the temporal comparisons; consequently, the assignment of samples to individual 50-year intervals, particularly below the unsupported-²¹⁰Pb dating horizons, should be regarded as approximate.

### DNA extraction and contamination control

DNA was extracted from 122 sediment samples using the DNeasy PowerMax Soil Kit (QIAGEN), following the manufacturer’s instructions, with a final elution volume of 1.2 mL. All extractions were performed in the dedicated clean DNA laboratory.

Potential contamination was monitored using seven subsampling blanks, eight extraction blanks and no-template PCR controls. Eight no-template PCR controls were included for each marker in each of the 11 sequencing libraries, resulting in 88 PCR controls per marker. DNA concentrations were measured using the Qubit High Sensitivity assay, and DNA extracts were stored at −20 °C until amplification.

### Metabarcoding library preparation and sequencing

Two metabarcoding markers were used to recover eukaryotic DNA from the sedimentary archives: a fragment of approximately 313 bp of the mitochondrial cytochrome c oxidase subunit I gene (COI), amplified using Leray-XT primers (Wangensteen et al., 2018), and an approximately 110-bp fragment of the V7 region of the nuclear 18S rRNA gene, amplified using 18S_allshort primers (Guardiola et al., 2015; Table S4).

Amplicon libraries were prepared using a twin-tagging dual-indexing strategy (Bohmann et al., 2022; Kircher et al., 2012). Identical 8-bp barcode sequences were attached to forward and reverse primers to minimise tag-switching artefacts (Schnell et al., 2015). Eight independent PCR replicates were performed per sample and marker to improve detection consistency and reduce stochastic amplification effects.

PCR reactions were performed in 40-µl volumes containing 4 µl of sedimentary DNA extract, 20 µl of AmpliTaq Gold 360 Master Mix (Thermo Fisher Scientific), 11.68 µl of laboratory water, 0.32 µl of BSA (20 mg ml⁻¹; Fisher Scientific) and 2 µl of each primer (5 nmol ml⁻¹). For COI, thermal cycling consisted of an initial denaturation at 95 °C for 10 min, followed by 35 cycles of 94 °C for 1 min, 45 °C for 1 min and 72 °C for 1 min, with a final extension at 72 °C for 5 min. For 18S, thermal cycling consisted of an initial denaturation at 95 °C for 10 min, followed by 45 cycles of 95 °C for 30 s, 45 °C for 30 s and 72 °C for 30 s, with a final extension at 72 °C for 5 min.

PCR products were purified using the MinElute PCR Purification Kit (QIAGEN) and quantified using Qubit. Eleven sequencing libraries were prepared using the TruSeq PCR-Free protocol (Illumina) and sequenced on an Illumina NovaSeq 6000 platform using paired-end sequencing (2 × 250 bp for COI; 2 × 150 bp for 18S). Negative controls were included without normalisation owing to their low DNA yield.

### Sequence processing and taxonomic assignment

Raw paired-end reads were demultiplexed using Cutadapt v2.3 (Martin, 2011), based on sample-specific dual barcodes and combined tag–primer sequences of approximately 36 bp. A maximum error rate of 0.1 was allowed, with insertions and deletions disallowed. Sense and antisense orientations were separated using the --pair-adapters and --pair-filter=both options and processed independently. Twin tags were designed with a minimum Hamming distance of three.

Amplicon sequence variants (ASVs) were inferred using DADA2 v1.12 (Callahan et al., 2016). Reads were quality-filtered using maxEE = 1 and truncQ = 2. Forward and reverse reads were truncated to 250 and 230 bp, respectively, for COI, and to 120 bp for both directions for 18S. Paired reads were merged with a minimum overlap of 18 bp, and chimeras were removed using removeBimeraDenovo. ASVs inferred from antisense reads were reverse-complemented and combined with those recovered from sense reads.

ASV read counts of three or fewer within individual samples were set to zero and ASVs represented by only one or two reads in a single PCR replicate across the entire dataset were removed as singletons or doubletons. To reduce potential contamination, the maximum read count observed for each ASV among negative controls was subtracted from its read count in each sample. ASVs were retained in the complete metabarcoding dataset regardless of taxonomic resolution, whereas species-level analyses were restricted to sequences meeting the assignment criteria described below.

Initial taxonomic assignment was performed using BLAST+ v2.14.1 (Camacho et al., 2009) against the NCBI nucleotide database (nt) downloaded in March 2024, retaining up to 300 alignments per query. Results were parsed using the custom R script ParseTaxonomy (doi:10.5281/zenodo.4671710). Taxonomic resolution was determined from sequence identity, query coverage and a lowest common ancestor approach. Initial species-level assignments required ≥97% sequence identity and ≥85% query coverage for COI, and 100% identity and ≥99% coverage for 18S. Ambiguous assignments and environmental labels were excluded.

For species-level analyses, ASVs initially assigned to Metazoa and Plantae were reassessed using the online NCBI BLAST interface against the nucleotide (nt) database in August 2026. Final species-level assignments required ≥97% identity and ≥95% query coverage for COI, and 100% identity and ≥99% coverage for 18S. ASVs that did not meet these thresholds, yielded ambiguous species-level matches or were reassigned to other taxa were excluded from species-level analyses. Validated taxonomic assignments were linked to the ASV table using unique ASV identifiers. Taxonomic nomenclature was harmonised using NCBI Taxonomy, and assignments labelled Viridiplantae were grouped under Plantae for graphical summaries.

All downstream data processing, statistical analyses and visualisations were performed in R (R Core Team, 2025).

### Sequencing-depth standardisation and sampling completeness

The influence of sequencing depth on ASV richness was assessed separately for COI and 18S. Each PCR replicate was randomly subsampled to 10,000 reads for COI and 20,000 reads for 18S. ASV richness before and after rarefaction was compared using Spearman’s rank correlation. The relationship between original sequencing depth and ASV richness after rarefaction was assessed using linear regression.

Sampling completeness was evaluated separately for each marker using sample-size-based and coverage-based rarefaction and extrapolation analyses implemented in iNEXT (Hsieh et al., 2016), with 95% confidence intervals. These analyses were based on the original, unrarefied ASV read abundances for individual PCR replicates. Both assessments were used as sensitivity checks of sequencing-depth standardisation and sampling completeness rather than as criteria for excluding samples from subsequent analyses.

### Species status classification and historical occurrence reconstruction

All Metazoa and Plantae represented by ASVs meeting the final species-level assignment criteria were considered for species status classification, comprising 202 species. Species status was assessed manually using biodiversity databases, non-indigenous species databases and published literature, with separate classifications for the northeastern Atlantic and western Mediterranean regions, focusing on the biogeographic context of the Iberian coastal sites.

Species were classified as native (NAT), non-indigenous (NIS) or cryptogenic (CRY). The cryptogenic category was used when native or introduced status could not be confidently established from the available evidence. Regional classifications were retained separately to account for differences in species status between the Atlantic and Mediterranean. Six combinations were observed: NAT in both regions, NIS in both regions, CRY in both regions, CRY-Med/NAT-Atl, NIS-Med/NAT-Atl and NAT-Med/NIS-Atl. For site-level analyses, each species was assigned the classification corresponding to the region of the sediment core, resulting in three analytical categories (NAT, NIS and CRY).

Species status classifications for all 202 species, together with ASV sequences and the supporting bibliographic sources for each assignment, are provided in Supplementary Data S1. Classifications were linked to ASVs through their final validated species-level assignments.

Historical occurrence patterns were reconstructed by linking validated species-level ASV detections to the estimated ages of the corresponding sediment horizons. COI and 18S were analysed separately. A species was considered detected in a sediment sample when at least one retained ASV assigned to that species was recovered in any of its PCR replicates after sequence filtering and contamination control. The number of positive PCR replicates was recorded for each species detection as a descriptive measure of detection consistency across PCR replicates. Occurrence plots were organised by site and marker, with point size indicating the number of positive PCR replicates, point shape representing taxonomic kingdom and colour indicating species status.

The earliest sedimentary detection was interpreted as the earliest recovered occurrence within the sampled archive, rather than as an estimate of the timing of introduction or establishment (Byers et al., 2015). Detections at different depths were not assumed to indicate uninterrupted local presence between sampled horizons, and non-detection was not interpreted as evidence of historical absence.

### Putative haplotype networks and temporal visualisation

Within each species–marker combination, distinct retained ASVs were treated as putative haplotypes to characterise within-species sequence variation and its observed geographic distribution (Estensmo et al., 2021; Skelton et al., 2023). These putative haplotypes represent unique metabarcoding sequences and were not independently validated as population-genetic haplotypes. Accumulated networks combined detections across all sampled sediment horizons without retaining their temporal ordering.

Sequences were aligned using AlignSeqs in DECIPHER v3.6.0 (Wright, 2016). Pairwise sequence distances were calculated using dist.dna in the R package *ape,* with pairwise deletion of missing positions (Paradis et al., 2004). Minimum spanning trees were constructed using mst in *igraph* for species represented by multiple ASVs; single-ASV species were displayed as single nodes. In accumulated networks, ASV read counts were summed by study site and displayed as site-coloured pie charts. Node sizes represented total read abundance, scaled within each network, and sectors indicated the relative contribution of each site. For the comparative networks (Fig. 2), sequence divergence was expressed as the percentage of differing aligned nucleotides, and edge widths followed a common scale across the selected networks.

Accumulated network structure was characterised using putative haplotype richness (number of distinct ASVs), dominant haplotype detection share, proportions of shared and site-specific haplotypes, and the mean number of sites per haplotype. Dominant haplotype detection share was defined as the number of sediment samples containing the most frequently detected ASV divided by the sum of sediment-sample detections across all ASVs assigned to that species, without counting PCR replicates as independent samples. Shared haplotypes were ASVs detected at two or more study sites, whereas site-specific haplotypes were detected at only one site within the sampled dataset; both proportions were calculated relative to total putative haplotype richness. Metrics were summarised separately for COI and 18S, and comparisons among species status categories were descriptive.

For selected representative taxa, temporal networks were reconstructed using common, approximately 50-year intervals based on estimated sediment ages. The accumulated network topology and node coordinates were held constant across temporal panels. Within each interval, pie-chart sectors represented the proportions of positive sediment samples contributed by each site, counting each sample only once per ASV. Node sizes reflected the number of positive sediment samples for each ASV, using a common scale within each species–marker temporal series. ASVs not detected in a given interval remained visible as small, unfilled nodes, and connections between ASVs detected in the same interval were highlighted. Each panel reported the number of detected ASVs and distinct sediment samples positive for the focal species. Only intervals with at least one detection of that species were displayed.

Network visualisations described recovered sequence variation and patterns of sedimentary detection, rather than population genetic diversity, dispersal, evolutionary replacement or continuous population persistence. Differences in marker resolution, sampling intensity and molecular detection were considered when interpreting these patterns.

### Temporal turnover and putative haplotype richness

Temporal changes in within-species ASV composition and richness were quantified separately for COI and 18S using presence–absence data grouped into approximately 50-year intervals. The main compositional analysis focused on 18S, with COI analysed as a complementary dataset. Comparisons between species status categories were restricted to native (NAT) and non-indigenous (NIS) taxa, excluding cryptogenic species (CRY).

For each species, site and period, putative haplotype composition was defined as the set of distinct ASVs detected in at least one sediment sample. All possible pairs of periods in which the same species was detected at a given site were compared using Jaccard dissimilarity, calculated as (1-|A\cap B|/|A\cup B|), where (A) and (B) are the ASV sets detected in each period (Legendre, 2014). Dissimilarity ranges from 0 for identical sets to 1 when no ASVs are shared. Species-level values were averaged by site, species status and temporal separation between periods, with each species–period comparison contributing equally. Point sizes in the resulting visualisations represented the number of comparisons contributing to each mean.

An additional analysis characterised changes between consecutive periods with detections of each species within a site, which were not necessarily adjacent 50-year intervals. The union of ASVs detected across each pair was partitioned into variants detected in both periods, only in the earlier period or only in the later period. Component proportions were calculated relative to the total number of distinct ASVs in the union and averaged across species within each site, species status and temporal transition, giving equal weight to each species.

Putative haplotype richness was calculated as the number of distinct ASVs detected per species, site and period. Richness was compared between NAT and NIS species within the same site and period using two-sided, unpaired Wilcoxon rank-sum tests when each category contained at least three species. Tests used the normal approximation, and P values were adjusted across eligible site–period comparisons using the Benjamini–Hochberg false discovery rate procedure. Analyses were conducted separately for COI and 18S.

These analyses quantified differences in recovered sedimentary ASV composition and richness. Non-detection was not interpreted as biological absence, and temporal differences were not taken as direct evidence of population loss, colonisation or changes in population genetic diversity.

## Results

### Sediment chronology and sequencing recovery

Sediment accumulation rates and chronological coverage varied considerably among the eight cores (Table S2; Fig. S1). Recent accumulation rates ranged from 0.10 cm yr⁻¹ at STO and CSD to 0.63 cm yr⁻¹ at COL and HIO, with intermediate values at DEE (0.11 cm yr⁻¹), CMB (0.15 cm yr⁻¹), HIT (0.42 cm yr⁻¹) and CCO (0.53 cm yr⁻¹). Basal ages extrapolated from the recent ²¹⁰Pb-derived accumulation rates ranged from approximately 1134 CE at STO to 1953 CE at COL. Chronological estimates were particularly uncertain for STO and CSD, where only two unsupported ²¹⁰Pb observations contributed to the regressions, while accumulation-rate uncertainty was also substantial at CMB and HIO.

AMS ¹⁴C determinations from deeper sections of five cores yielded calibrated ages spanning the first millennium BCE to the fifteenth century CE (Table S3; Fig. S2). In four cores (DEE, CSD, HIT and HIO), these ages were older than the corresponding basal ages extrapolated from recent ²¹⁰Pb-derived accumulation rates, whereas the younger calibrated interval for CMB approached its extrapolated basal estimate. These comparisons were interpreted cautiously because the radiocarbon determinations dated carbon of uncertain depositional origin and were obtained below the unsupported-²¹⁰Pb horizons.

Metabarcoding generated 1,377,919,936 raw reads across COI and 18S, including sediment samples and negative controls. Following amplification and sequence filtering, 114 of the 122 extracted sediment samples were retained for 18S and 106 for COI. The final 18S dataset comprised 424,652 ASVs and 748,479,852 reads across 829 PCR replicates, whereas the COI dataset comprised 270,136 ASVs and 399,543,516 reads across 832 PCR replicates. Mean sequencing depth per retained PCR replicate was 902,871 reads for 18S and 480,221 reads for COI.

Rarefaction to 20,000 reads for 18S and 10,000 reads for COI reduced mean ASV richness by 23% and 43%, respectively, while retaining strong correlations with richness before rarefaction (Spearman’s ρ = 0.954 for 18S and 0.856 for COI; Fig. S3A). Original sequencing depth explained a small proportion of the variation in rarefied richness (18S: R² = 0.076; COI: R² = 0.048; Fig. S3B). Sample-size-based and coverage-based rarefaction and extrapolation curves indicated variation in sampling completeness among PCR replicates (Fig. S3C,D).

### Historical species detections and species status classifications across sedimentary archives

Sedimentary DNA detections revealed contrasting temporal and geographic patterns among taxa classified as non-indigenous, cryptogenic or assigned different species status categories in the northeastern Atlantic and western Mediterranean (Fig. S4). Several taxa were recovered from sediment horizons assigned historical ages, including taxa whose documented occurrence histories in the study regions remain uncertain.

Among the selected metazoans, the ascidians *Ascidia ahodori* and *Ecteinascidia turbinata*, classified as non-indigenous in the study region, were detected in sediment horizons assigned ages before 1500 CE at Cádiz site CSD. The green crab *Carcinus aestuarii*, classified as non-indigenous in the Atlantic region, was also recovered from a Cádiz sediment horizon assigned an age before 1600 CE.

Plant detections included *Oryza sativa*, recovered from Ebro Delta sediments assigned ages before 1400 CE, as well as from other study sites, with the earliest Cádiz detection assigned an age before 1500 CE (Fig. S5). The taxon assigned to *Suaeda taxifolia* was detected at three southern sites (CSD, HIT and HIO), including a CSD horizon assigned to the thirteenth century.

COI and 18S recovered partly distinct sets of taxa, and detections varied in their distribution across sediment horizons (Fig. S4). Additional plant taxa were recovered from multiple sites and historical periods (Fig. S5). These records document the recovery of species-assigned sequences from sediment horizons of different estimated ages, without establishing the timing of species introduction, establishment or continuous local presence.

### Accumulated haplotype diversity and geographic sharing

Accumulated networks revealed substantial variation in putative haplotype richness, dominance and geographic sharing across species–marker combinations (Fig. 2). This variation was also evident within each species status category, with no consistent descriptive separation among native (NAT), non-indigenous (NIS) and cryptogenic (CRY) taxa across the complete dataset (Fig. S6; Table S5).

COI networks illustrated contrasting patterns of within-species sequence variation. The native bivalve *Cerastoderma edule* contained five putative haplotypes, 20% of which were shared among sites, whereas the non-indigenous ascidian *A. ahodori* contained four haplotypes, with 50% shared. The cryptogenic-in-the-Mediterranean gastropod *Peringia ulvae* showed the highest richness among the representative networks, with 84 COI haplotypes, a dominant-haplotype detection share of 0.04 and 15% shared among sites. Other taxa displayed more restricted geographic sharing, including *Styela plicata*, for which none of the three recovered COI haplotypes was detected at multiple sites.

The 18S networks also varied in richness and geographic distribution. The cryptogenic ascidian *Botrylloides israeliense* contained 30 putative haplotypes, of which 63% were shared among sites. All three 18S haplotypes recovered for the non-indigenous sea anemone *Diadumene lineata* were detected at multiple sites. In contrast, *C. aestuarii*, classified as non-indigenous in the Atlantic, was represented by a single 18S haplotype recovered across four sites. The native *C. edule* contained two 18S haplotypes, both detected at multiple sites, compared with five COI haplotypes.

Across all species–marker combinations, median COI haplotype richness was 3.0 for NAT, 3.5 for NIS and 5.0 for CRY taxa, with median proportions of geographically shared haplotypes of 0.14, 0.32 and 0.20, respectively (Fig. S6; Table S5). For 18S, median richness was two haplotypes in all three categories, whereas median proportions of shared haplotypes were 0.23, 0.17 and 0 for NAT, NIS and CRY taxa, respectively. These descriptive summaries indicated considerable variation among species within each status category, rather than a consistent pattern distinguishing native from non-indigenous taxa.

**Fig. 2:**
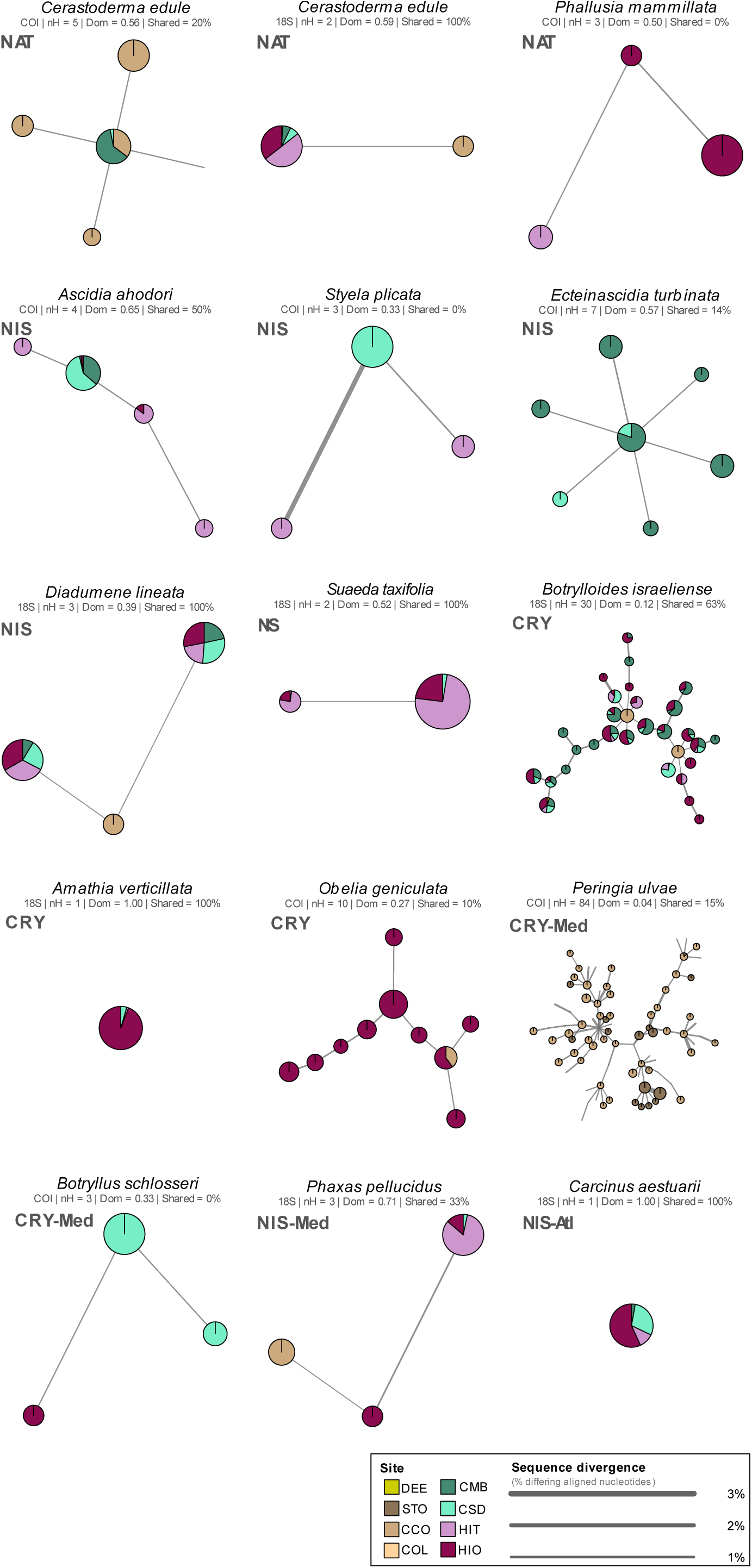
Accumulated putative haplotype diversity and geographic sharing across representative taxa. Minimum spanning trees depict distinct metabarcoding ASVs treated as putative haplotypes for selected species with contrasting regional status classifications. Node size represents total ASV read abundance, scaled independently within each network, and pie-chart sectors indicate the relative contributions of study sites. Edges connect ASVs according to minimum spanning tree topology, with widths scaled to pairwise sequence divergence (% differing aligned nucleotide positions) using a common scale across networks. Network labels indicate the metabarcoding marker, putative haplotype richness (nH), dominant-haplotype detection share (Dom) and percentage of haplotypes detected at two or more study sites (Shared). NAT, native; NIS, non-indigenous; CRY, cryptogenic; Atl, northeastern Atlantic; Med, western Mediterranean.

### Historical changes in haplotype composition

Temporal reconstructions revealed differences in the richness, identity and geographic distribution of putative haplotypes recovered across approximately 50-year intervals (Fig. 3). The accumulated networks comprised variants that were not necessarily detected together within individual periods, and the magnitude of temporal variation differed among the four representative species.

**Fig. 3:**
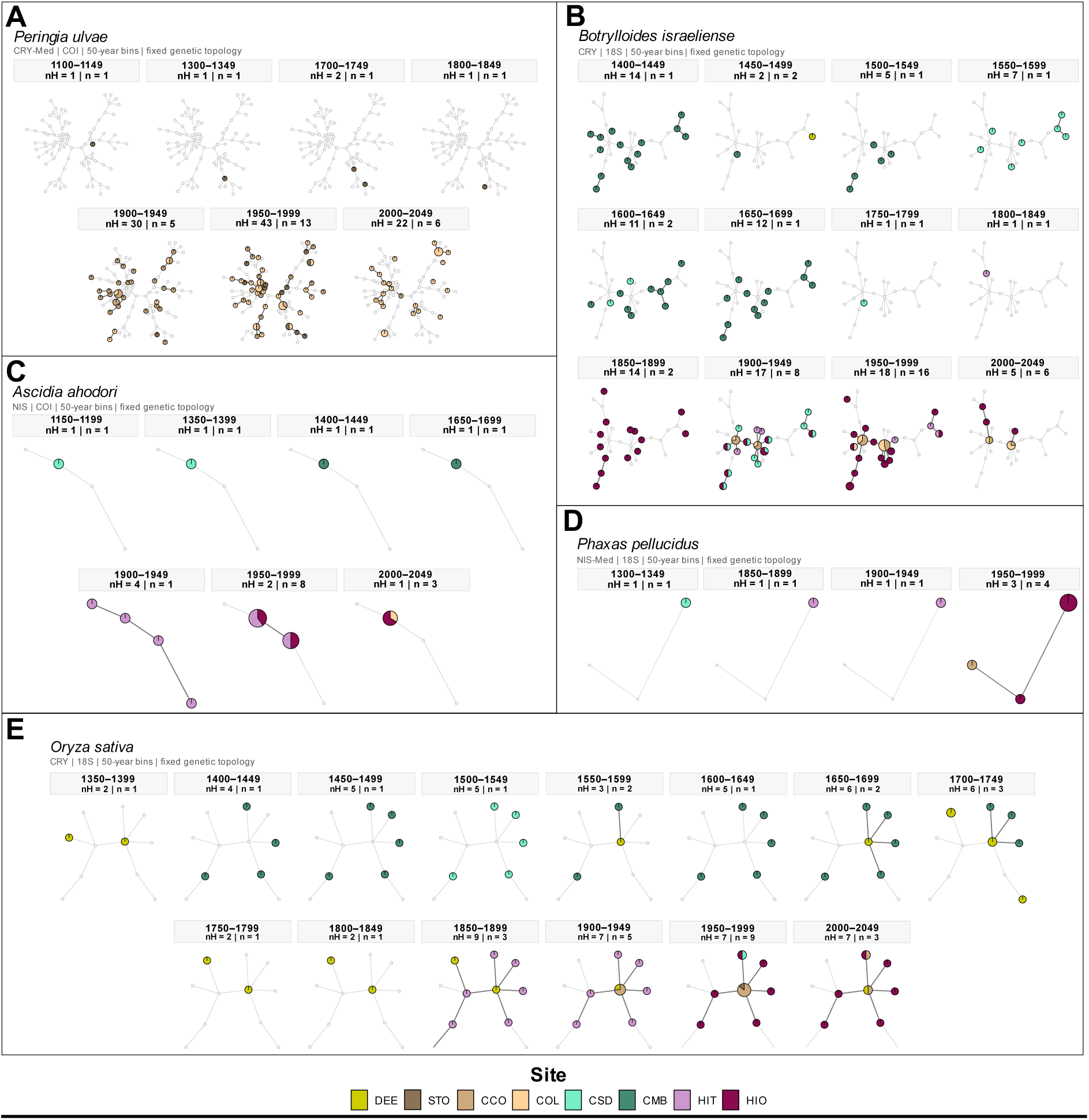
Temporal variation in putative haplotype composition and geographic distribution. Minimum spanning tree networks show putative haplotypes recovered across approximately 50-year intervals for Peringia ulvae (COI), Botrylloides israeliense (18S), Ascidia ahodori (COI), Phaxas pellucidus (18S) and Oryza sativa (18S). Network topology and node positions remain fixed across temporal panels for each species. Coloured nodes indicate ASVs detected within an interval, with node size proportional to the number of positive sediment samples and pie-chart sectors indicating their relative contributions from study sites. Small, unfilled nodes represent ASVs not detected in the corresponding interval. Panel labels indicate the number of detected putative haplotypes (nH) and distinct sediment samples positive for the focal species (n). Only intervals containing detections are displayed.

*P. ulvae* showed pronounced temporal variation in detected richness, with only one or a few haplotypes recovered in the earliest represented intervals, compared with 30 in 1900–1949, 43 in 1950–1999 and 22 in the interval beginning in 2000. Thus, no individual interval contained all 84 haplotypes represented in its accumulated network. *B. israeliense* (18S) also showed substantial temporal differences, with 14 haplotypes detected in 1850–1899, 17 in 1900–1949, 18 in 1950–1999 and five in the interval beginning in 2000. For both species, the identities of detected variants and their distribution among study sites differed across periods.

Simpler temporal configurations were observed in *A. ahodori* (COI) and *P. pellucidus* (18S). Earlier detections of *A. ahodori* were dominated by a single haplotype, followed by four detected haplotypes in 1900–1949, two in 1950–1999 and one in the interval beginning in 2000. For *P. pellucidus*, one haplotype was recovered in each of the 1300–1349, 1850–1899 and 1900–1949 intervals, increasing to three in 1950–1999.

These temporal reconstructions showed that accumulated network structure combined distinct configurations of recovered ASVs across sedimentary intervals. Differences in detected richness and composition were not interpreted as direct evidence of population-level haplotype gains, losses or replacement.

### Temporal turnover in within-species sequence composition

Within-species ASV composition varied across historical periods, with substantial differences in mean Jaccard dissimilarity among sites and species status categories in the 18S dataset (Fig. 4A; Table S6). Mean dissimilarity was lowest at CCO (NAT: 0.101; NIS: 0.139) and comparatively low at COL (NAT: 0.210; NIS: 0.216). Higher values were observed at other sites, with non-indigenous taxa showing greater mean dissimilarity than native taxa at CMB (0.525 versus 0.435), CSD (0.593 versus 0.266), DEE (0.507 versus 0.204), HIT (0.525 versus 0.356) and STO (0.667 versus 0.439). These differences were descriptive and varied in magnitude among sites. No consistent relationship was apparent between Jaccard dissimilarity and increasing temporal separation between periods.

**Fig. 4:**
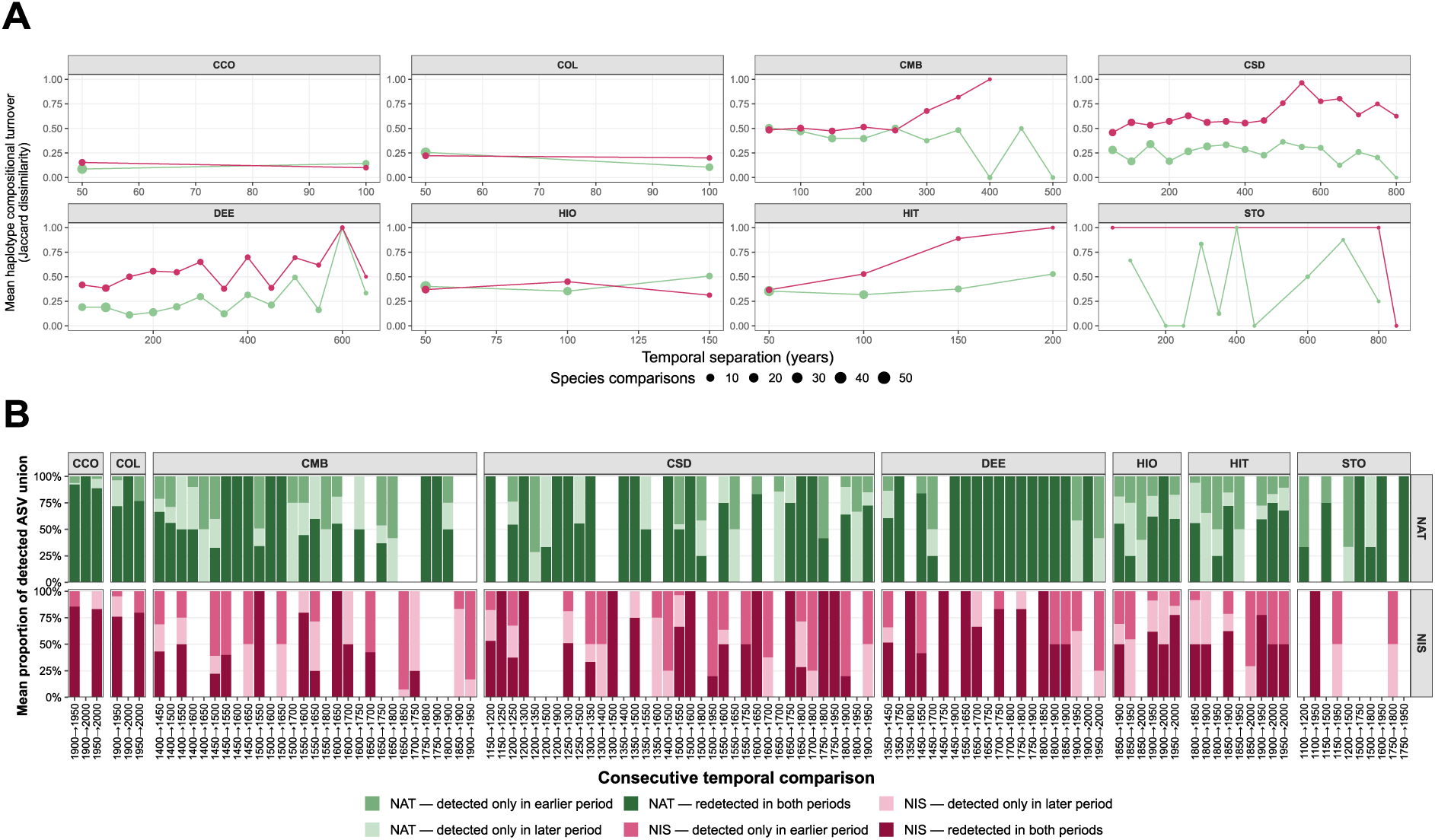
Temporal turnover in within-species ASV composition. Analyses are based on 18S ASVs treated as putative haplotypes. a, Mean Jaccard dissimilarity between approximately 50-year periods by study site and species status (native, NAT; non-indigenous, NIS), plotted against temporal separation. Values range from 0 (identical detected ASV sets) to 1 (no shared ASVs). Point size indicates the number of species–period comparisons contributing to each mean; lines represent descriptive trends. b, Mean proportions of ASVs detected in both periods, only in the earlier period or only in the later period, for consecutive periods with detections of each species within a site. These periods are not necessarily adjacent 50-year intervals. Proportions are calculated relative to the union of ASVs detected across each pair and averaged across species with equal weight. Only species detected in both compared periods are included.

Comparisons between consecutive periods with detections of each species showed that compositional differences reflected varying proportions of ASVs detected in both periods or in only one of the two periods (Fig. 4B; Table S6). The mean proportion detected in both periods was highest at CCO and COL, reaching 0.932 and 0.787 for native taxa and 0.844 and 0.810 for non-indigenous taxa, respectively. Lower proportions were observed at CMB (NAT: 0.469; NIS: 0.362). The relative contributions of ASVs detected only in the earlier or later period also varied among sites and species status categories.

Putative haplotype richness comparisons provided a complementary assessment (Fig. S7; Table S7). Across 36 eligible 18S site–period comparisons, differences between native and non-indigenous taxa varied in direction and magnitude. The largest observed difference occurred at HIT in 1850–1899, where mean richness was 4.429 ASVs per non-indigenous species and 2.037 per native species (unadjusted P = 0.002; FDR-adjusted P = 0.077). No comparison remained statistically significant after false discovery rate correction. For COI, no site–period combination met the minimum requirement of three species per status category.

Overall, temporal turnover differed among sites and species, whereas the richness analyses did not identify statistically supported differences between native and non-indigenous taxa after correction for multiple testing.

## Discussion

Contemporary genetic observations provide only a partial perspective on how within-species sequence variation has been distributed through time (Casas-Marce et al., 2017). Historical genetic records can reveal changes in genetic diversity and structure that may not be resolved from contemporary samples alone (Habel et al., 2014; Ramakrishnan & Hadly, 2009). Here, sedimentary DNA from eight Iberian coastal wetlands showed that accumulated networks combined putative haplotypes recovered from different historical intervals and geographic locations (Figs. 3 and 4). Temporally resolved networks revealed differences in the identity, richness and geographic representation of detected variants, while analyses of the broader 18S dataset identified heterogeneous compositional turnover among sites and taxa (Fig. 4). The accumulated diversity of an environmental archive therefore does not necessarily represent the sequence composition recovered within any individual historical period.

This distinction was particularly evident in *P. ulvae*. Its accumulated COI network contained 84 putative haplotypes, yet no individual temporal interval contained all these variants. Detected richness reached 30 haplotypes in 1900–1949 and 43 in 1950–1999, whereas only a few were recovered in the earliest represented intervals (Figs. 3 and 4). The identities of detected variants and their geographic representation also differed among periods. However, sedimentary coverage varied among sites: the STO core extended further into the past than those from CCO and COL (Fig. S1; Table S2). Consequently, the contribution of additional sites to later intervals cannot be interpreted as evidence of recent dispersal. Rather, the networks demonstrate that the accumulated set of recovered variants represents different combinations of sedimentary detections across space and time. Comparable temporal differences occurred in *B. israeliense*, *A. ahodori* and *P. pellucidus,* despite their contrasting accumulated network structures.

Temporal networks have been used to represent genetic variation in serially sampled material (Prost & Anderson, 2011), while sedimentary DNA has revealed historical changes in within-species sequence composition, including shifts in mitochondrial haplotype dominance in lake-dwelling rotifers (Epp et al., 2010). More recent studies have further demonstrated the recovery of intra-specific sequence variation from sedimentary ancient DNA (Kyalo-Omamo et al., 2023; Lammers et al., 2024). Here, combining temporal networks with detections from multiple coastal sites allowed us to examine how the recovered variants and their geographic representation differed among sedimentary intervals. These patterns would be obscured if all detections were considered only as a single accumulated network.

The broader 18S analysis indicated that temporal differences in recovered ASV composition were not restricted to the taxa selected for network visualisation. Jaccard dissimilarity varied among sites and species status categories, with comparatively low values at CCO and COL and greater compositional differences at several other sites (Fig. 4; Table S6). No consistent increase in dissimilarity with temporal separation was apparent. Moreover, comparisons between consecutive periods with detections were not always comparisons between adjacent 50-year intervals. These patterns therefore describe heterogeneous changes in sedimentary sequence recovery rather than a uniform trajectory of genetic differentiation.

Accumulated haplotype richness, dominance and geographic sharing varied widely within both native and non-indigenous taxa, without a consistent separation between the two categories. Although accumulated network richness, dominance and geographic sharing varied among species, their distributions did not show a consistent descriptive separation by species status across COI and 18S (Fig. 2; Fig. S6). Likewise, within-site and period comparisons of putative haplotype richness showed no statistically significant differences between native and non-indigenous taxa after correction for multiple testing (Fig. S7; Table S7). This does not demonstrate equivalence between categories, particularly given differences in sampling coverage and marker resolution. More importantly, similar numbers of recovered variants need not imply similar sequence composition or temporal histories. Richness and compositional turnover therefore provide complementary, rather than interchangeable, descriptions of within-species variation.

The absence of a consistent pattern associated with species status should also be considered in the context of biological introductions. Introduced populations can experience contrasting genetic outcomes, including reductions in diversity following founder events or the maintenance and accumulation of variation through multiple introductions and admixture (Kolbe et al., 2004; Lavergne & Molofsky, 2007; Rius & Darling, 2014). Consequently, no single genetic signature is expected across all introduced taxa (Dlugosch & Parker, 2008; Estoup et al., 2016; Roman & Darling, 2007). However, the sedimentary patterns observed here cannot be attributed to particular introduction histories, founder events or demographic processes. Regional species status identifies whether a taxon is considered native, non-indigenous or cryptogenic in a given biogeographic context; it does not directly characterise the processes responsible for its sequence variation. Cryptogenic taxa likewise represent uncertainty in biogeographic classification rather than a biologically homogeneous group (Carlton, 1996).

Historical species detections provide an additional perspective on regional status classifications and documented occurrence histories. Sedimentary sequences assigned to the ascidians *A. ahodori* and *E. turbinata* were recovered from Cádiz horizons with estimated ages predating 1500 CE, while *C. aestuarii*, classified here as non-indigenous in the Atlantic region, was detected in a horizon assigned an age before 1600 CE (Fig. S4). The latter species is primarily associated with the Mediterranean, although its historical distribution near the Atlantic–Mediterranean transition in southwestern Iberia remains uncertain (Carlton & Cohen, 2003; Darling, 2011). Plant detections also raise questions about historical occurrence. Sequences assigned to *Oryza sativa* were recovered from Ebro Delta horizons with estimated ages predating 1400 CE (Fig. S5 and 4). Although rice cultivation was established in the Iberian Peninsula during the twelfth century (Sharma, 2010), historical accounts place the earliest reported attempts to cultivate rice in the Ebro Delta in the fifteenth century (Sabarís, 1958), with extensive cultivation developing following nineteenth-century irrigation works (Dobby, 1936). Although these sedimentary detections are consistent with a potentially longer history of rice occurrence in the delta, they cannot establish the timing or location of cultivation. Sequences assigned to *S. taxifolia* were detected at three southern sites (Fig. S5 and 3), including a CSD horizon assigned to the thirteenth century, despite the species’ documented distribution being centred on coastal California. The estimated ages of these detections raise questions about the documented histories of several taxa, but do not establish when they first appeared or became established in the study regions. Taxonomic uncertainty, sedimentary DNA transport and preservation, and the extrapolated age–depth models are particularly important when evaluating such unexpected assignments. Independent taxonomic and chronological corroboration would be needed before interpreting them as confirmed historical range records.

Marker resolution is a further constraint on the biological interpretation of recovered sequence variation. For example, *C. edule* was represented by five COI variants but only two 18S variants (Fig. 2). COI and 18S differ in sequence conservation, amplification characteristics and their ability to distinguish variation within species (Wangensteen et al., 2018). Previous metabarcoding studies have demonstrated the recovery of within-species sequence variation while also identifying limitations associated with amplification, sequencing and sequence inference (Elbrecht et al., 2018; Serrana & Watanabe, 2023; Tsuji et al., 2019). The ASVs examined here are therefore considered putative haplotypes, not independently validated population-genetic haplotypes. In particular, a single recovered 18S variant does not establish genetic homogeneity, and identical ASVs recovered at different sites do not independently demonstrate contemporary or historical connectivity between populations.

Sedimentary processes and chronological uncertainty impose additional limitations. DNA recovered from a sediment horizon may reflect production, transport, deposition, preservation and molecular recovery processes operating over different spatial and temporal scales (Capo et al., 2021; Crump, 2021). Accordingly, non-detection of a variant cannot establish its historical absence, while repeated recovery does not necessarily indicate continuous local persistence. Unequal numbers of sediment samples among sites and periods can also influence detected haplotype richness and geographic sharing. Although sequencing-depth assessments characterised the effects of read depth and sampling completeness (Fig. S3), they did not standardise the number of sediment samples contributing to each temporal interval. Furthermore, the age–depth models relied partly on extrapolating recent ²¹⁰Pb-derived accumulation rates beyond their directly supported horizons, and chronological uncertainty was not propagated into the 50-year comparisons (Fig. S1; Table S2). Radiocarbon determinations provided independent age information but were not incorporated as depositional constraints because the dated carbon fractions were not independently characterised (Table S3). These limitations are particularly important when interpreting detections assigned to older intervals or comparing the apparent timing of changes between cores.

These limitations do not diminish the broader potential of sedimentary DNA for reconstructing biodiversity change across ecosystems. Dated sedimentary records have revealed long-term ecological changes in freshwater lakes (Ficetola et al., 2018), shifts in alpine plant diversity associated with climate and land use (Garcés-Pastor et al., 2022), and historical changes in marine communities (Rivera Rosas et al., 2024). Recovering distinguishable variants within species adds a potentially informative dimension to such records, particularly where contemporary observations or historical specimens provide limited temporal coverage. Historical collections have already demonstrated the value of directly comparing genetic material across periods (Wandeler et al., 2007). Combining sedimentary archives with well-characterised reference sequences, independent chronological constraints and contemporary or historical population-genetic data could help determine which changes in recovered ASV composition correspond to biological variation and which arise from differences in environmental DNA recovery.

Sedimentary DNA records can reveal differences in within-species sequence composition that are not apparent from species occurrences or time-aggregated haplotype networks alone. Across the wetlands examined here, the variants recovered within individual periods differed in identity, richness and geographic representation, although these patterns were not uniform among taxa or sites. Accounting for sampling effort, marker resolution, sedimentary processes and chronological uncertainty will be essential for determining how much of this variation reflects biological change. With these constraints in mind, temporally resolved metabarcoding offers a way to extend historical biodiversity reconstructions beyond species-level occurrence records.

## Acknowledgments

This work was supported by the TEMPOINVASIONS grants TED2021-132228B-C21 and TED2021-132228B-C22, both funded by MCIN/AEI/10.13039/501100011033 and the European Union (“NextGenerationEU”/PRTR). We are also grateful to the Spanish Ministry of Science, Innovation and Universities for providing financial support with the BlueDNA PID2023-146307OB-C22 grant (MICIU/AEI/https://doi.org/10.13039/501100011033 and FEDER, UE). Pawel Gaca and Madeleine Cobbold are thanked for conducting ^210^Pb dating analyses. We thank Antonio Moreno Muñoz, for his assistance and collaboration during fieldwork in Cádiz and Huelva. We thank Maria Rigat for her contribution to preliminary species status assignments for taxa from the Santoña and Ebro Delta datasets. Marc Rius and Xavier Turon are members of the research group SGR2021-00405 and Sandra Nogué is member of the research group SGR 01333, both funded by the Generalitat de Catalunya. Luke E. Holman is supported by the European Research Council through the SeaChange project (Grant agreement No. 856488).

## Author contributions

Elena Baños, Marc Rius, Xavier Turon, Erik de Boer, Jesús Zarcero, Xavier Benito, Enrique González Ortegón, and Sandra Nogué conducted the fieldwork and collected the samples. Dating analyses were carried out by Andrew B. Cundy and Elena Baños and interpreted by Andrew B. Cundy, Sandra Nogué and Elena Baños. Elena Baños performed the laboratory work and conducted the bioinformatic analyses of metabarcoding datasets. Data analyses and statistical interpretation was carried out by Elena Baños. Elena Baños wrote the manuscript with major contributions from Sandra Nogué and Marc Rius. All authors contributed to manuscript revision and approved the final version.

## Data Availability Statement

Raw COI and 18S metabarcoding sequence data are available through the NCBI Sequence Read Archive under BioProject PRJNA1535453. The processed datasets, metadata, supporting tables and R scripts used to reproduce the statistical analyses and figures are publicly available on GitHub and archived on Zenodo under DOI 10.5281/zenodo.22880848.

## Supplementary tables

**Table S1:** Sampling locations and core characteristics for sedimentary DNA (*seda*DNA) analyses. Geographic coordinates, site names, and total core depths are reported for each sampling location.

| **Core ID** | **Site** | **Latitude** | **Longitude** | **Core Depth (cm)** |
|---|---|---|---|---|
| DEE | Ebro Delta Encanyissada | 40.653108 N | 0.685247 E | 70 |
| STO | Santoña | 43.446528 N | -3.466719 W | 90 |
| CCO | Cicero | 43.422497 N | -3.481806 W | 50 |
| COL | Colindres | 43.400247 N | -3.455278 W | 50 |
| CSD | Cadiz Salina Dolores | 36.464889 N | -6.223167 W | 90 |
| CMB | Cadiz Militar Base | 36.499139 N | -6.193056 W | 90 |
| HIT | Huelva Isla Saltes Tinto River | 37.181778 N | -6.937831 W | 90 |
| HIO | Huelva Isla Saltes Ocean | 37.180806 N | -6.938719 W | 90 |

**Table S2:** Recent sediment accumulation rates estimated from excess ^210^Pb profiles using the Constant Flux–Constant Sedimentation (CF:CS) model. Sediment accumulation rates and their 95% uncertainty ranges were derived from linear regressions of ln-transformed excess ^210^Pb activity against sediment depth. The unsupported-^210^Pb dating horizon indicates the deepest interval at which excess ^210^Pb remained distinguishable from supported background activity. Basal ages are extrapolations assuming that the recent CF:CS accumulation rate remained constant below the ^210^Pb dating horizon and should not be interpreted as independently dated ages. † Estimates for STO and CSD are poorly constrained because their regressions contained only two unsupported-^210^Pb observations.

| Core ID | Site | Unsupported<sup>210</sup>Pb dating horizon (cm) | CF:CS SAR (cm yr<sup>-1</sup>) | 95% range (cm yr<sup>-1</sup>) | Core base (cm) | Extrapolated basal age assuming constant CF:CS SAR (CE) |
|---|---|---|---|---|---|---|
| DEE | Ebro Delta | 15 | 0.11 | 0.09 - 0.15 | 70 | ~1384 |
| STO | Santoña | 10 | 0.1 | 0.1† | 89 | ~1134 |
| CCO | Cicero | 48 | 0.53 | 0.42 - 0.73 | 49 | ~1939 |
| COL | Colindres | 49 | 0.63 | 0.50 - 0.87 | 49 | ~1953 |
| CSD | Cadiz Salina Dolores | 3 | 0.1 | 0.1† | 89 | ~1135 |
| CMB | Cadiz Militar Base | 15.5 | 0.15 | 0.08 - 0.58 | 89 | ~1419 |
| HIT | Huelva Tinto River | 40 | 0.42 | 0.3019 - 0.6850 | 89 | ~1812 |
| HIO | Huelva Saltes Island Ocean | 65 | 0.63 | 0.417 - 1.302 | 89 | ~1884 |

**Table S3:** AMS radiocarbon determinations and calibrated probability ranges for intertidal sediment samples. Conventional radiocarbon ages were calibrated using the IntCal20 Northern Hemisphere atmospheric calibration curve. Reported ranges encompass 95.4% of the calibrated probability distribution; the probability associated with each discontinuous interval is shown in parentheses. No marine-reservoir or old-carbon correction was applied.

| **Core ID** | **Depth (cm)** | **Conventional<sup>14</sup>C age (yr BP)** | **1σ error (yr)** | **Calibration curve** | **Calibrated age, 95.4% HPD (cal BCE/CE)** |
|---|---|---|---|---|---|
| DEE | 70 | 1570 | 30 | IntCal20 | 424–564 CE (95.6%) |
| CSD | 85 | 2735 | 35 | IntCal20 | 975–954 BCE (4.0%);<br>933–809 BCE (91.5%) |
| CMB | 85 | 580 | 30 | IntCal20 | 1305–1364 CE (64.3%);<br>1383–1419 CE (31.2%) |
| HIT | 85 | 955 | 30 | IntCal20 | 1027–1159 CE (95.5%) |
| HIO | 85 | 1095 | 30 | IntCal20 | 889–995 CE (90.2%);<br>1005–1018 CE (5.4%) |

**Table S4:**
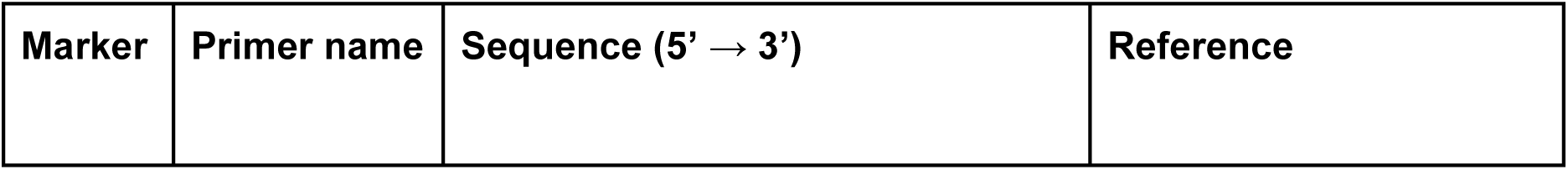
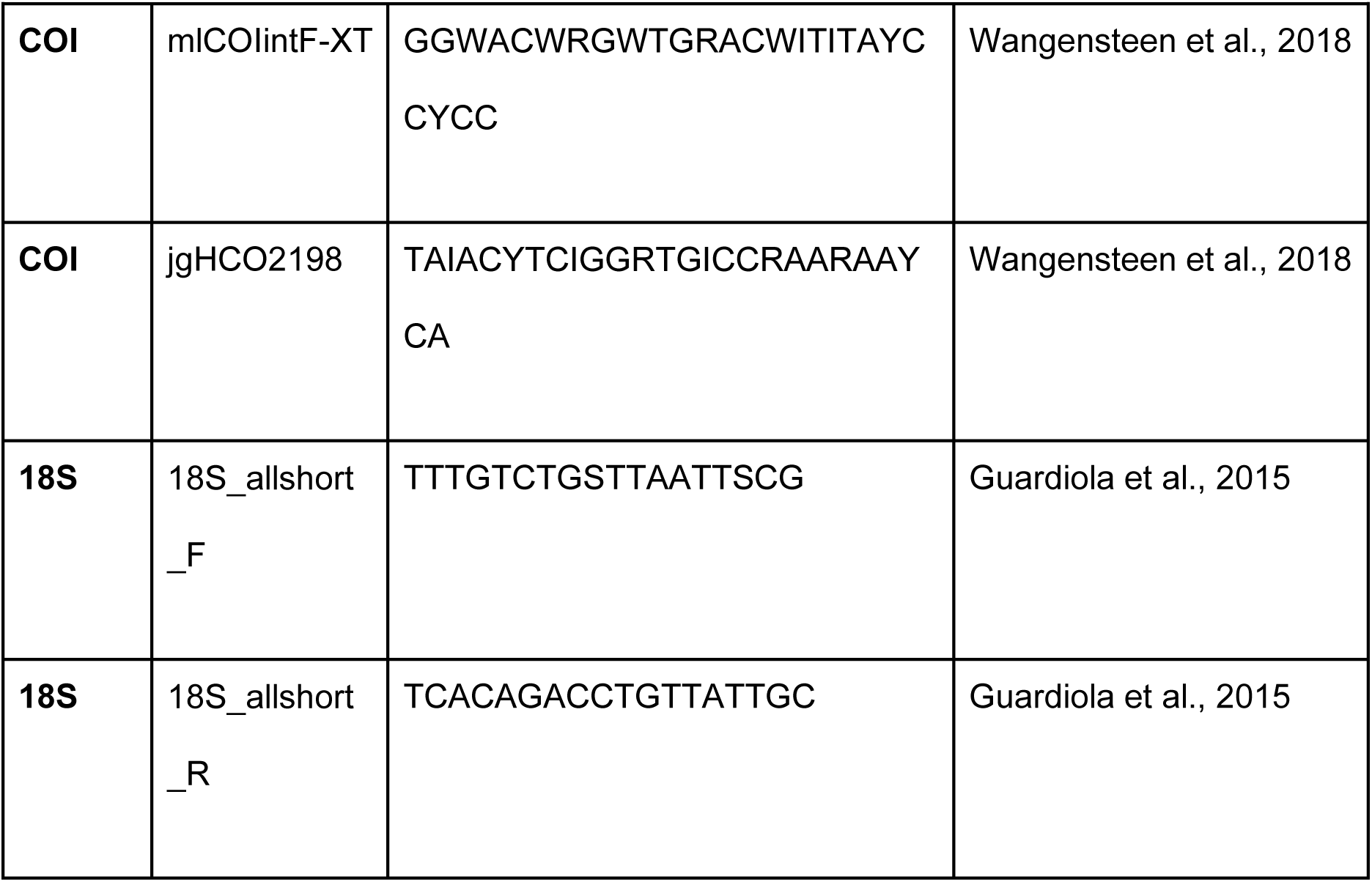
Primer sequences (5′–3′) used to amplify the COI (Leray-XT) and 18S (V7 region) markers in this study. Primers were dual-indexed using mirrored 8-bp tags to minimise tag-jumping artefacts.

**Table S5:** Summary of haplotype-structure metrics by species status and metabarcoding marker. For each status and marker, n indicates the number of species. H, haplotype richness (number of putative haplotypes/ASVs detected per species); D, dominant haplotype frequency; SH, proportion of shared haplotypes (detected at ≥2 study sites); SSH, proportion of site-specific haplotypes (detected at a single site); S/H, mean number of sites occupied per haplotype. Med., median. Mean and median values are reported for each metric. NAT, native; NIS, non-indigenous; CRY, cryptogenic.

| **Status** | **Marker** | **n** | **H**<br>**median** | **H**<br>**mean** | **D**<br>**median** | **D**<br>**mean** | **SH**<br>**median** | **SH**<br>**mean** | **SSH**<br>**median** | **SSH**<br>**mean** | **S/H**<br>**median** | **S/H**<br>**mean** |
|---|---|---|---|---|---|---|---|---|---|---|---|---|
| NAT | COI | 11 | 3.00 | 4.91 | 0.50 | 0.59 | 0.14 | 0.21 | 0.86 | 0.79 | 1.32 | 1.28 |
| NAT | 18S | 102 | 2.00 | 4.35 | 0.61 | 0.66 | 0.23 | 0.38 | 0.78 | 0.62 | 1.33 | 1.63 |
| NIS | COI | 4 | 3.50 | 3.75 | 0.61 | 0.64 | 0.32 | 0.41 | 0.68 | 0.59 | 1.57 | 1.60 |
| NIS | 18S | 37 | 2.00 | 6.22 | 0.52 | 0.60 | 0.17 | 0.30 | 0.83 | 0.70 | 1.27 | 1.58 |
| NIS-Atl | 18S | 1 | 1.00 | 1.00 | 1.00 | 1.00 | 1.00 | 1.00 | 0.00 | 0.00 | 4.00 | 4.00 |
| NIS-Med | COI | 1 | 3.00 | 3.00 | 0.50 | 0.50 | 0.33 | 0.33 | 0.67 | 0.67 | 1.33 | 1.33 |
| NIS-Med | 18S | 3 | 3.00 | 2.33 | 0.71 | 0.74 | 0.33 | 0.44 | 0.67 | 0.56 | 1.67 | 1.56 |
| CRY | COI | 25 | 5.00 | 8.96 | 0.47 | 0.49 | 0.20 | 0.24 | 0.80 | 0.76 | 1.29 | 1.39 |
| CRY | 18S | 25 | 2.00 | 3.56 | 0.75 | 0.71 | 0.00 | 0.37 | 1.00 | 0.63 | 1.00 | 1.56 |
| CRY-Med | COI | 2 | 43.50 | 43.50 | 0.19 | 0.19 | 0.08 | 0.08 | 0.92 | 0.92 | 1.08 | 1.08 |
| CRY-Med | 18S | 4 | 4.00 | 4.25 | 0.33 | 0.42 | 0.45 | 0.44 | 0.55 | 0.56 | 1.45 | 1.51 |

**Table S6:** Summary of temporal within-species haplotype compositional turnover in the 18S dataset across study sites and status. Nsp, number of species contributing to all temporal comparisons; Ncomp, number of species-level temporal comparisons; Lag min and Lag max, minimum and maximum temporal separation between compared 50-year periods, respectively; Jmean, mean Jaccard dissimilarity across all species-level temporal comparisons; *Nconsec*, number of comparisons between consecutive periods in which a species was detected; Shared, mean proportion of putative haplotypes redetected in both periods; Earlier-only, mean proportion detected only in the earlier period; Later-only, mean proportion detected only in the later period; and Jconsec, mean Jaccard dissimilarity for consecutive detected-period comparisons. Component proportions were first averaged across consecutive comparisons within each species and subsequently across species, giving equal weight to each species. Consecutive detected periods were not necessarily adjacent 50-year periods when a species was not detected in an intervening period. Earlier-only and later-only represent differences in detection and should not be interpreted as biological loss, extinction, gain or arrival. NAT, native; NIS, non-indigenous species.

| **Site** | **Status** | **n** | **Ncomp** | **Lag min** | **Lag max** | **J mean** | **Shared** | **Earlier-only** | **Later-only** | **Jconsec** |
|---|---|---|---|---|---|---|---|---|---|---|
| CCO | NAT | 32 | 58 | 50 | 100 | 0.101 | 0.932 | 0.042 | 0.026 | 0.068 |
|  | NIS | 8 | 18 | 50 | 100 | 0.139 | 0.844 | 0.094 | 0.063 | 0.156 |
| COL | NAT | 27 | 65 | 50 | 100 | 0.210 | 0.787 | 0.110 | 0.103 | 0.213 |
|  | NIS | 7 | 17 | 50 | 100 | 0.216 | 0.810 | 0.095 | 0.095 | 0.190 |
| CMB | NAT | 22 | 119 | 50 | 500 | 0.435 | 0.469 | 0.258 | 0.273 | 0.531 |
|  | NIS | 7 | 68 | 50 | 400 | 0.525 | 0.362 | 0.350 | 0.289 | 0.638 |
| CSD | NAT | 13 | 206 | 50 | 800 | 0.266 | 0.465 | 0.197 | 0.338 | 0.535 |
|  | NIS | 7 | 149 | 50 | 800 | 0.593 | 0.530 | 0.304 | 0.167 | 0.470 |
| DEE | NAT | 17 | 199 | 50 | 650 | 0.204 | 0.726 | 0.114 | 0.160 | 0.274 |
|  | NIS | 6 | 100 | 50 | 650 | 0.507 | 0.684 | 0.217 | 0.098 | 0.316 |
| HIO | NAT | 32 | 94 | 50 | 150 | 0.400 | 0.564 | 0.195 | 0.241 | 0.436 |
|  | NIS | 12 | 34 | 50 | 150 | 0.386 | 0.578 | 0.177 | 0.245 | 0.422 |
| HIT | NAT | 28 | 101 | 50 | 200 | 0.356 | 0.558 | 0.208 | 0.234 | 0.442 |
|  | NIS | 9 | 20 | 50 | 200 | 0.525 | 0.495 | 0.330 | 0.175 | 0.505 |
| STO | NAT | 3 | 13 | 100 | 800 | 0.439 | 0.713 | 0.176 | 0.111 | 0.287 |
|  | NIS | 3 | 3 | 50 | 850 | 0.667 | 0.333 | 0.333 | 0.333 | 0.667 |

**Table S7:** Within-period comparisons of detected putative haplotype richness between native and non-indigenous taxa from the 18S dataset. Detected putative haplotype richness was defined as the number of distinct ASVs detected per species within each site and 50-year period. Only site–period combinations containing at least three native (NAT) and three non-indigenous (NIS) species were included in statistical comparisons. NAT n and NIS n indicate the number of species represented in each status category; NAT mean and NIS mean are the corresponding mean numbers of detected putative haplotypes per species; NAT median and NIS median are the corresponding medians; and Δmean is the difference in mean richness calculated as NIS − NAT. W is the Wilcoxon rank-sum test statistic, praw is the unadjusted p-value, and pFDR is the p-value adjusted for multiple comparisons using the Benjamini– Hochberg false discovery rate procedure. No COI site–period combination met the minimum representation criterion for statistical comparison; therefore, the table contains only eligible 18S comparisons. NAT, native; NIS, non-indigenous species.

| Site | Period | NAT n | NIS n | NAT mean | NIS mean | NAT median | NIS median | Δ mean | W | praw | pFDR |
|---|---|---|---|---|---|---|---|---|---|---|---|
| CCO | 1900–1949 | 38 | 8 | 1.158 | 1.250 | 1 | 1 | 0.092 | 173 | 0.309 | 0.873 |
|  | 1950–1999 | 41 | 15 | 1.098 | 1.000 | 1 | 1 | -0.098 | 292.5 | 0.404 | 0.873 |
|  | 2000–2049 | 17 | 6 | 1.294 | 1.333 | 1 | 1 | 0.039 | 58 | 0.526 | 0.873 |
| COL | 1900–1949 | 28 | 8 | 1.143 | 1.125 | 1 | 1 | -0.018 | 113.5 | 0.944 | 0.944 |
|  | 1950–1999 | 43 | 7 | 1.628 | 1.429 | 1 | 1 | -0.199 | 164 | 0.652 | 0.909 |
|  | 2000–2049 | 24 | 5 | 1.208 | 1.200 | 1 | 1 | -0.008 | 64 | 0.735 | 0.909 |
| CMB | 1400–1449 | 18 | 8 | 3.167 | 3.625 | 2 | 2.5 | 0.458 | 85.5 | 0.453 | 0.873 |
|  | 1450–1499 | 14 | 6 | 2.357 | 3.500 | 2 | 2 | 1.143 | 53.5 | 0.334 | 0.873 |
|  | 1500–1549 | 13 | 3 | 3.538 | 2.333 | 2 | 2 | -1.205 | 14.5 | 0.533 | 0.873 |
|  | 1550–1599 | 7 | 5 | 2.000 | 2.000 | 2 | 2 | 0.000 | 16.5 | 0.929 | 0.944 |
|  | 1600–1649 | 12 | 3 | 2.667 | 3.333 | 2 | 4 | 0.667 | 22 | 0.599 | 0.909 |
|  | 1650–1699 | 16 | 6 | 3.500 | 4.333 | 2 | 3 | 0.833 | 52.5 | 0.759 | 0.909 |
| CSD | 1200–1249 | 9 | 4 | 2.444 | 2.250 | 2 | 2 | -0.194 | 20 | 0.806 | 0.909 |
|  | 1250–1299 | 5 | 3 | 2.400 | 2.667 | 2 | 3 | 0.267 | 11 | 0.353 | 0.872 |
|  | 1300–1349 | 7 | 4 | 2.000 | 3.000 | 2 | 2 | 1.000 | 20 | 0.269 | 0.872 |
|  | 1500–1549 | 12 | 7 | 2.417 | 2.714 | 2 | 2 | 0.298 | 47 | 0.691 | 0.909 |
|  | 1550–1599 | 4 | 4 | 1.500 | 3.750 | 1.5 | 3 | 2.250 | 12 | 0.288 | 0.872 |
|  | 1600–1649 | 4 | 4 | 2.000 | 3.000 | 2 | 3 | 1.000 | 14 | 0.066 | 0.583 |
|  | 1800–1849 | 7 | 3 | 2.000 | 2.667 | 1 | 2 | 0.667 | 13 | 0.621 | 0.909 |
|  | 1900–1949 | 7 | 4 | 2.429 | 1.250 | 1 | 1 | -1.179 | 10.5 | 0.510 | 0.872 |
|  | 1950–1999 | 7 | 3 | 2.286 | 1.000 | 2 | 1 | -1.286 | 3 | 0.077 | 0.583 |
| DEE | 1350–1399 | 29 | 9 | 1.276 | 2.667 | 1 | 1 | 1.391 | 172 | 0.058 | 0.583 |
|  | 1450–1499 | 21 | 7 | 1.429 | 1.571 | 1 | 1 | 0.143 | 86.5 | 0.403 | 0.872 |
|  | 1700–1749 | 6 | 3 | 1.000 | 1.667 | 1 | 1 | 0.667 | 12 | 0.238 | 0.872 |
|  | 1750–1799 | 7 | 3 | 1.000 | 1.333 | 1 | 1 | 0.333 | 14 | 0.190 | 0.856 |
|  | 1800–1849 | 5 | 4 | 1.000 | 1.500 | 1 | 1 | 0.500 | 15 | 0.128 | 0.659 |
| HIO | 1850–1899 | 20 | 9 | 2.150 | 2.111 | 1.50 | 2 | -0.039 | 94.5 | 0.839 | 0.909 |
|  | 1900–1949 | 30 | 11 | 2.333 | 1.364 | 1.50 | 1 | -0.970 | 116.5 | 0.118 | 0.659 |
|  | 1950–1999 | 38 | 12 | 2.237 | 2.583 | 2 | 2 | 0.346 | 236 | 0.858 | 0.909 |
|  | 2000–2049 | 27 | 11 | 2.704 | 2.273 | 2 | 2 | -0.431 | 137.5 | 0.725 | 0.909 |
| HIT | 1800–1849 | 17 | 3 | 1.412 | 1.667 | 1 | 1 | 0.255 | 27.5 | 0.850 | 0.909 |
|  | 1850–1899 | 27 | 7 | 2.037 | 4.429 | 2 | 4 | 2.392 | 164.5 | 0.002 | 0.077 |
|  | 1900–1949 | 35 | 11 | 2.571 | 2.909 | 2 | 2 | 0.338 | 200 | 0.849 | 0.909 |
|  | 1950–1999 | 16 | 3 | 1.688 | 1.333 | 2 | 1 | -0.354 | 18 | 0.488 | 0.872 |
|  | 2000–2049 | 12 | 7 | 2.417 | 1.429 | 2 | 1 | -0.988 | 22 | 0.081 | 0.583 |
| STO | 1950–1999 | 20 | 4 | 1.250 | 1.000 | 1 | 1 | -0.250 | 34 | 0.458 | 0.872 |

## Supplementary figures

**Fig. S1:**
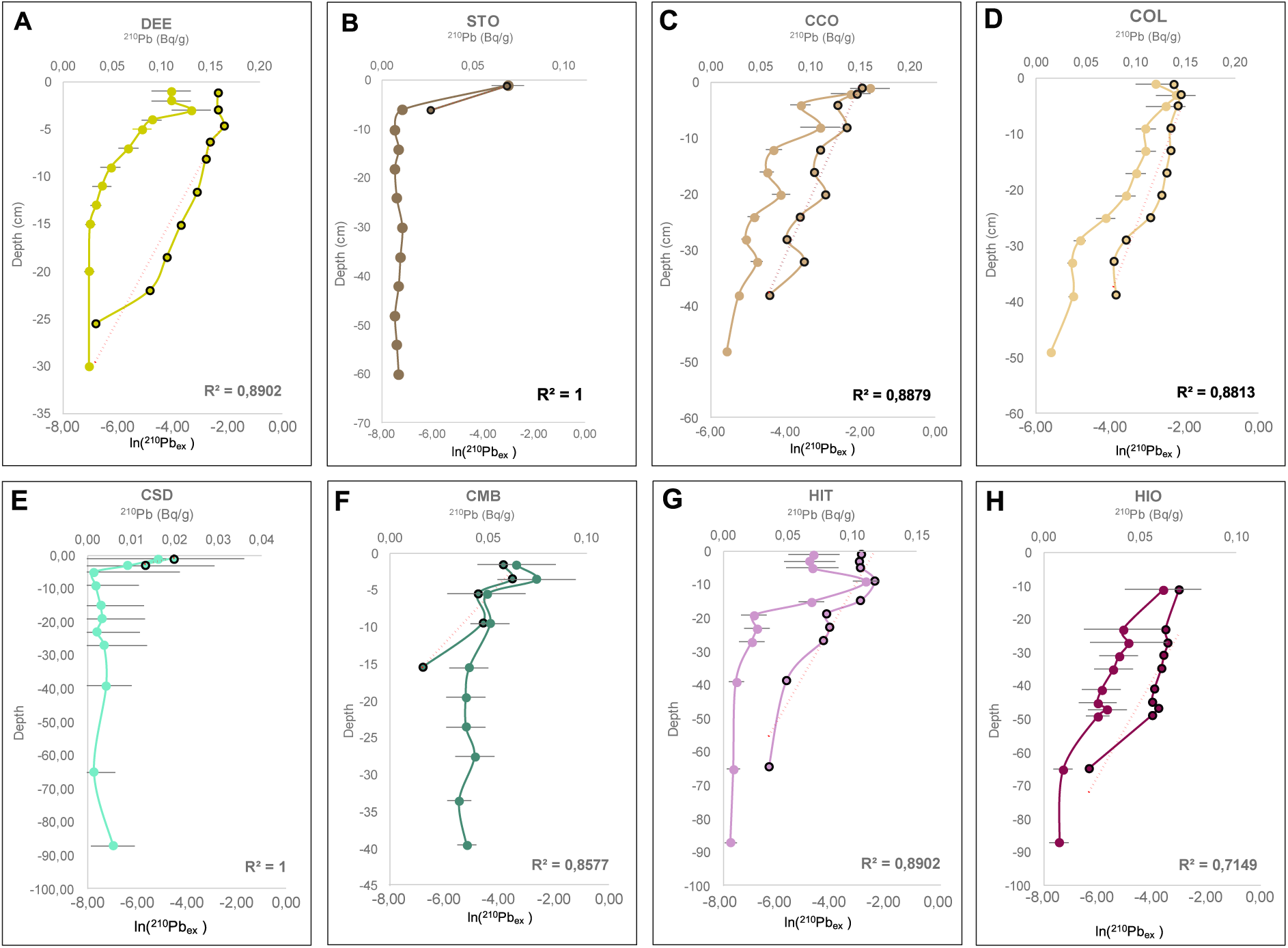
Excess ^210^Pb dating models for Iberian coastal sediment cores based on the Constant Flux–Constant Sedimentation (CF:CS) model. Plots show the relationship between the natural logarithm of unsupported ^210^Pb activity and sediment depth for each dated core. Linear regressions were used to estimate sediment accumulation rates (SAR), assuming constant sedimentation and radionuclide flux through time. Dashed lines indicate fitted regression models, and shaded areas represent the corresponding uncertainty intervals where available. Depths at which supported ^210^Pb activity was reached were used to define the limit of unsupported radionuclide inventories. Estimated sediment accumulation rates and basal age approximations are summarised in Table S2.

**Fig. S2:**
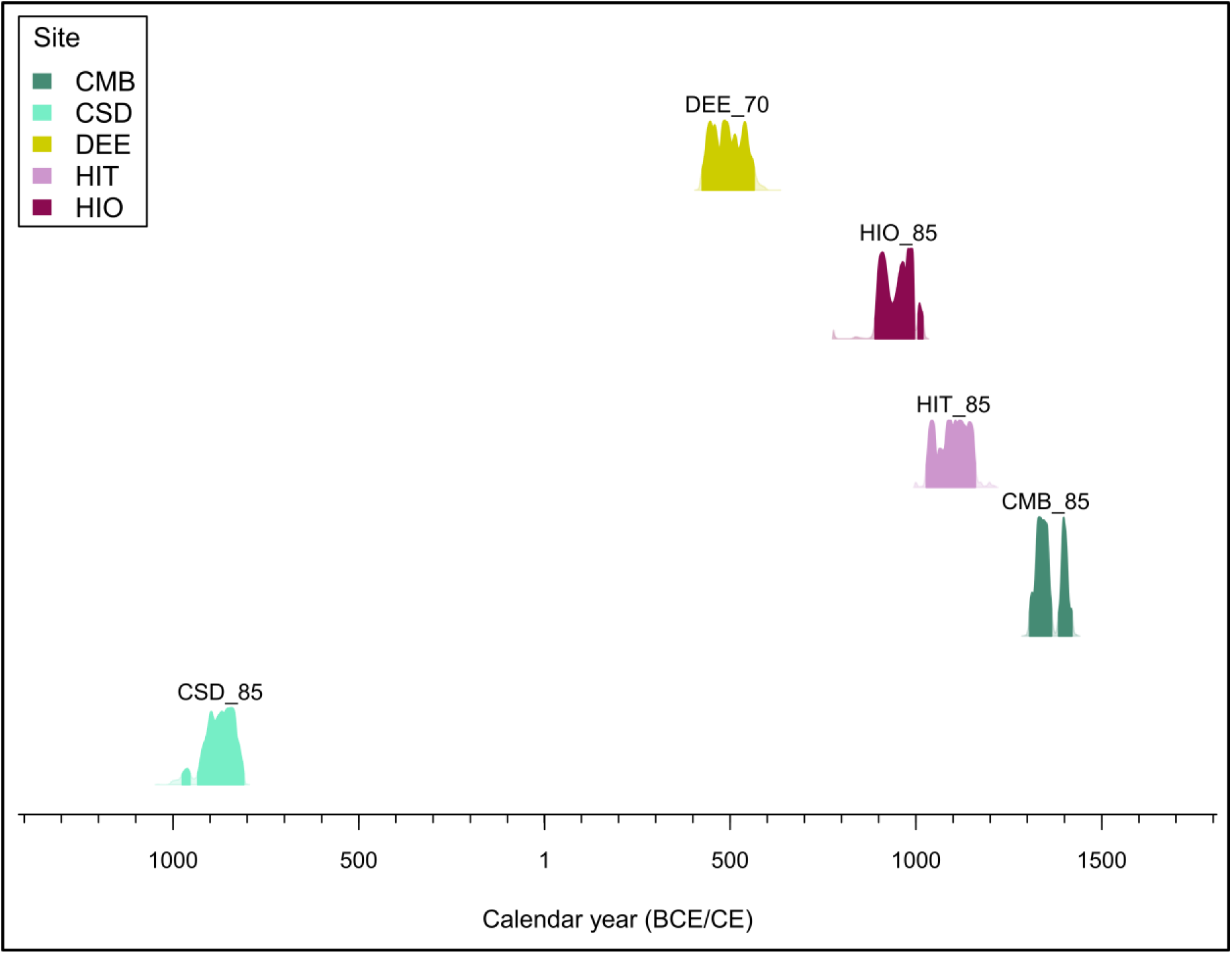
Calibrated AMS radiocarbon ages for intertidal sediment samples from five sediment cores. Conventional ^14^C ages were calibrated using the IntCal20 Northern Hemisphere atmospheric calibration curve. Coloured distributions represent the calibrated probability density for each sample, and horizontal bars indicate the intervals encompassing 95.4% of the calibrated probability distribution. Colours identify the corresponding sediment cores. No marine-reservoir or old-carbon correction was applied. The resulting age distributions represent the estimated ages of the dated carbon fractions and were not assumed a priori to correspond exactly to the depositional ages of the surrounding sediments. Conventional radiocarbon determinations and calibrated age ranges are summarised in Table S3.

**Fig. S3:**
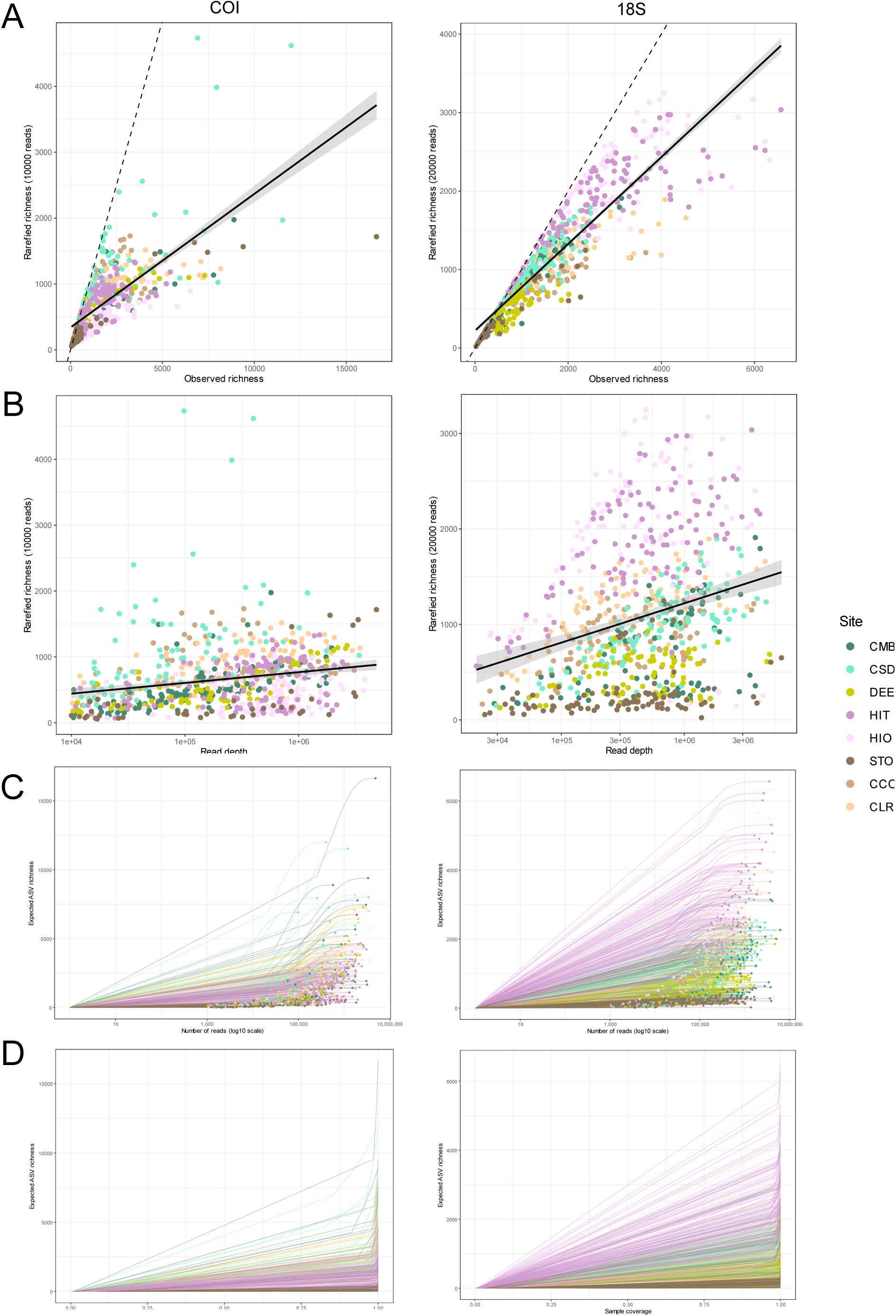
Evaluation of sequencing-depth standardisation and sampling completeness for the 18S and COI sedimentary DNA metabarcoding datasets. (A) Association between ASV richness before and after read-depth standardisation across PCR replicates. Libraries were rarefied to 20,000 reads for 18S and 10,000 reads for COI; the dashed line represents the 1:1 relationship between observed and rarefied richness. (B) Relationship between sequencing depth and ASV richness after rarefaction, used to evaluate residual effects of library size following standardisation. Lines represent fitted linear relationships with 95% confidence intervals. (C) Sample-size-based rarefaction and extrapolation curves generated with iNEXT for individual PCR replicates. Solid and dashed portions of the curves represent interpolation and extrapolation, respectively, with shaded areas indicating 95% confidence intervals. (D) Coverage-based rarefaction and extrapolation curves showing diversity estimates as a function of sample coverage, thereby allowing comparison of sampling completeness independently of differences in sequencing depth. Results are shown separately for the 18S and COI markers.

**Fig. S4:**
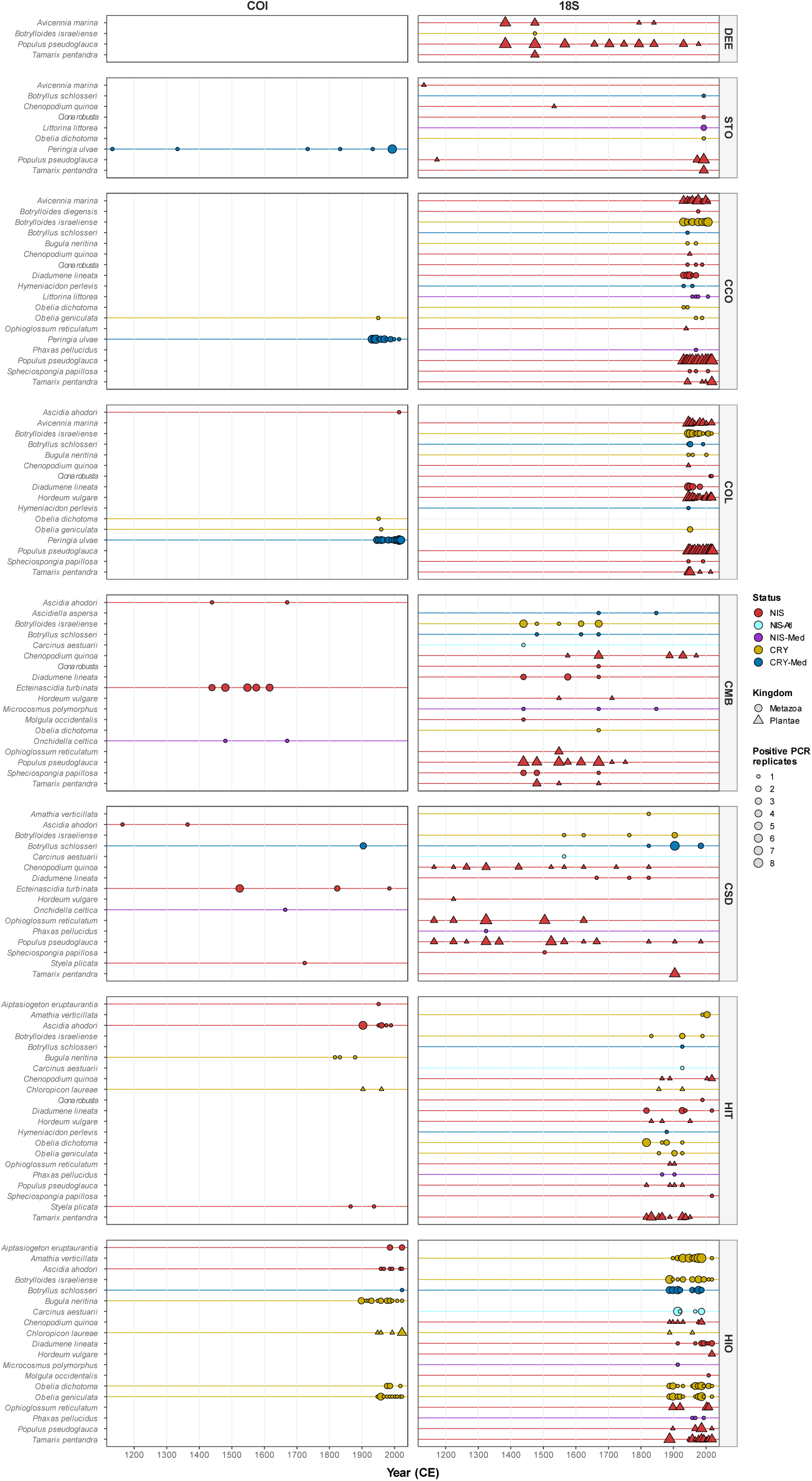
Historical sedimentary DNA detections of selected taxa with contrasting regional species status. Detections across eight coastal sediment cores are shown separately for COI (left) and 18S (right). Each point represents a taxon detected in a sediment sample assigned an estimated calendar age. Point size indicates the number of positive PCR replicates, shape distinguishes Metazoa (circles) from Plantae (triangles), and colour indicates species status: non-indigenous (NIS), cryptogenic (CRY), non-indigenous in the Atlantic (NIS-Atl), non-indigenous in the Mediterranean (NIS-Med) or cryptogenic in the Mediterranean (CRY-Med). Horizontal lines connect the earliest and latest sedimentary detections displayed for each taxon and do not imply continuous presence between them. Only selected taxa are shown. Sedimentary detections do not establish dates of introduction or establishment.

**Fig. S5:**
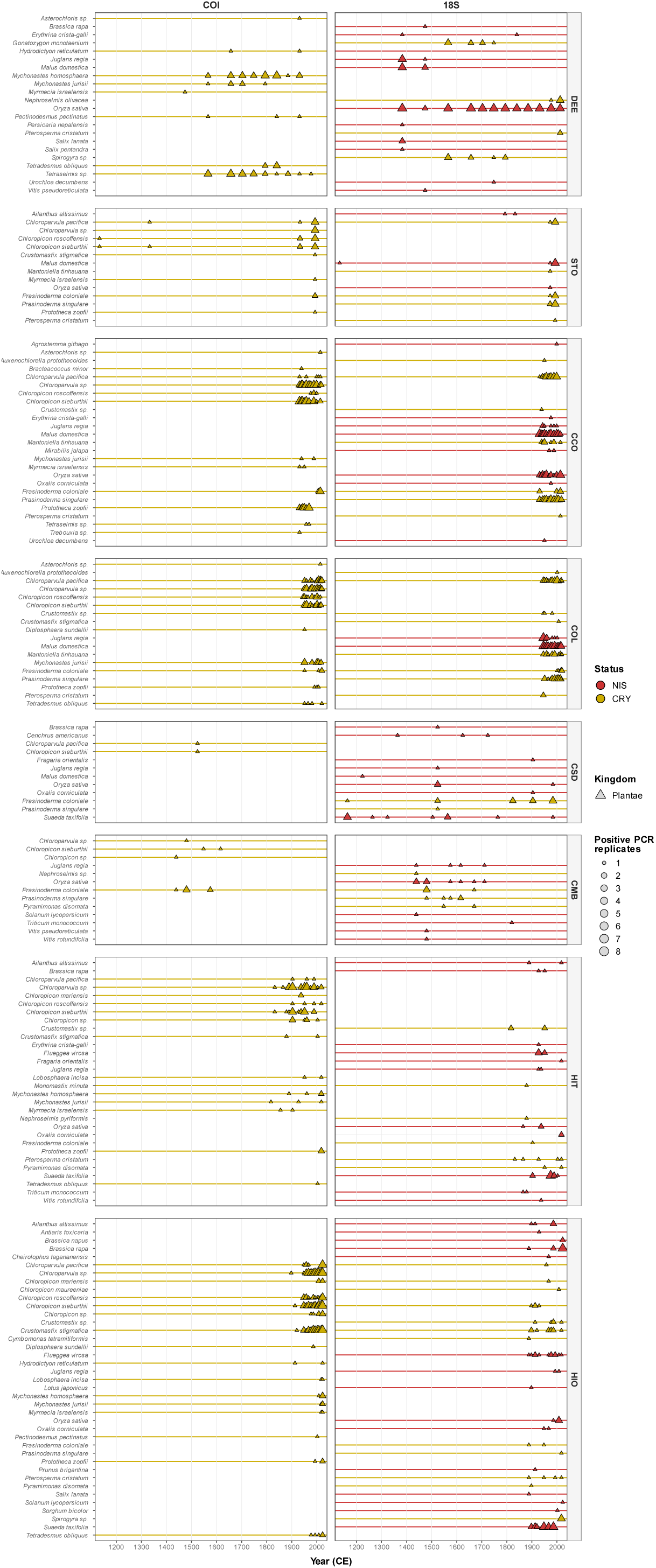
Long-term occurrence of additional plant taxa detected in coastal sedimentary DNA archives. Temporal distribution of plant taxa that were not selected for inclusion in the main-text occurrence timeline, shown separately for COI and 18S metabarcoding markers across the eight study sites. Each point represents detection of a taxon in a dated sediment horizon, with point size indicating the number of positive PCR replicates (1–8). Colours denote status: native (NAT), non-indigenous (NIS), native in the Mediterranean and non-indigenous in the Atlantic (NAT/NIS), non-indigenous in the Mediterranean and native in the Atlantic (NIS/NAT), cryptogenic (CRY), and cryptogenic in the Mediterranean and native in the Atlantic (CRY/NAT). Triangles indicate Plantae. Horizontal coloured lines connect the temporal range between the earliest and latest sedaDNA detections of each taxon within each site and marker. SedaDNA occurrence represents detection within the sedimentary archive and should not be interpreted as the timing of species introduction, establishment or local presence in the absence of detection.

**Fig. S6:**
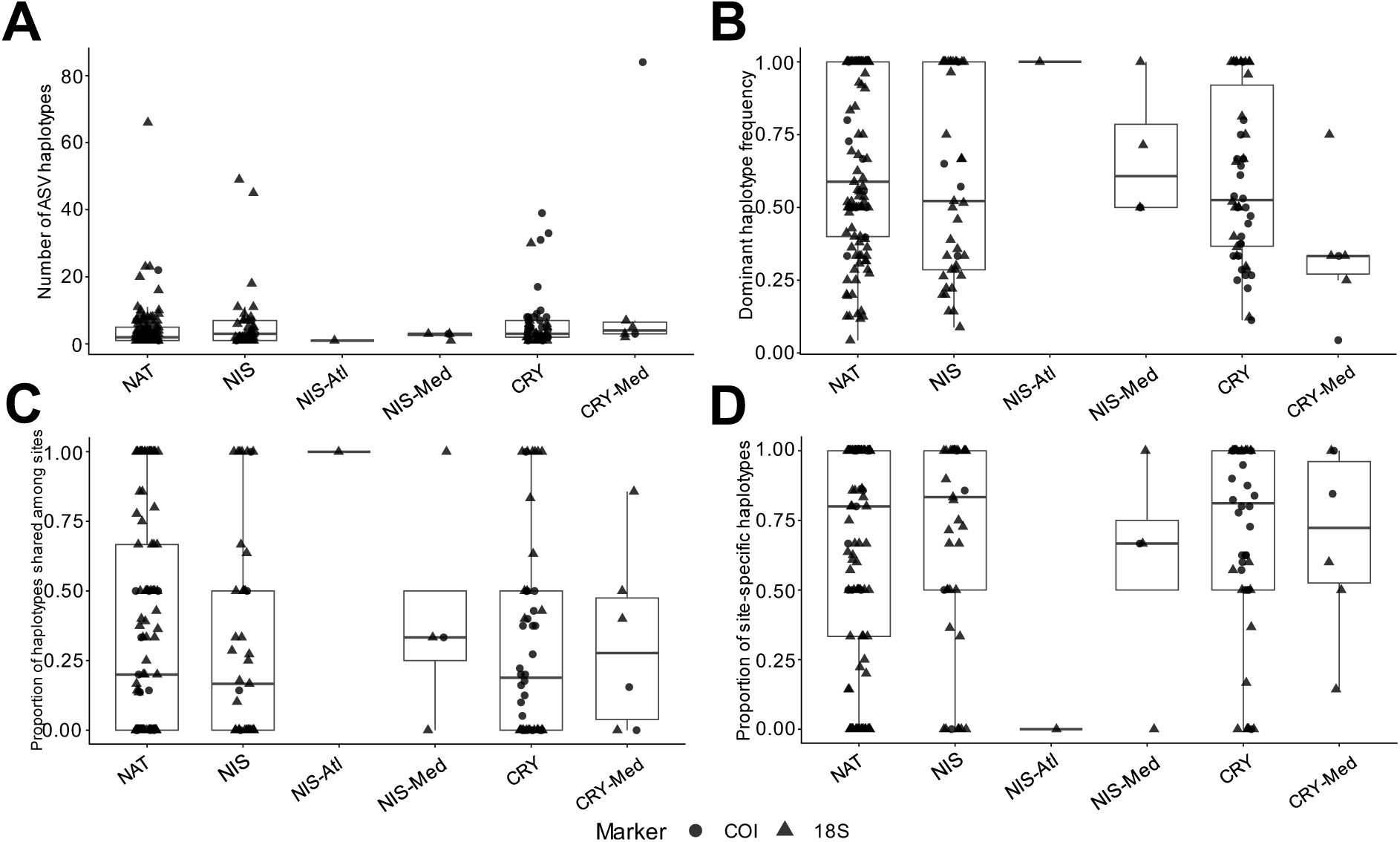
Quantitative variation in haplotype structure among status categories. Species-level haplotype metrics are shown separately for the 18S and COI markers and grouped according to status: native (NAT), non-indigenous (NIS), native in the Mediterranean and non-indigenous in the Atlantic (NIS-Atl), non-indigenous in the Mediterranean and native in the Atlantic (NIS-Med), cryptogenic (CRY), and cryptogenic in the Mediterranean and native in the Atlantic (CRY-Med). (A) Haplotype richness, expressed as the number of detected ASVs used as putative haplotypes; (B) dominant haplotype frequency, representing the relative contribution of the most frequently detected haplotype; (C) proportion of haplotypes shared among two or more study sites; and (D) proportion of haplotypes restricted to a single study site. Points represent individual species–marker combinations, while boxplots summarize their distribution within each status category. Metrics are shown separately by marker because haplotype resolution differs between 18S and COI.

**Fig. S7:**
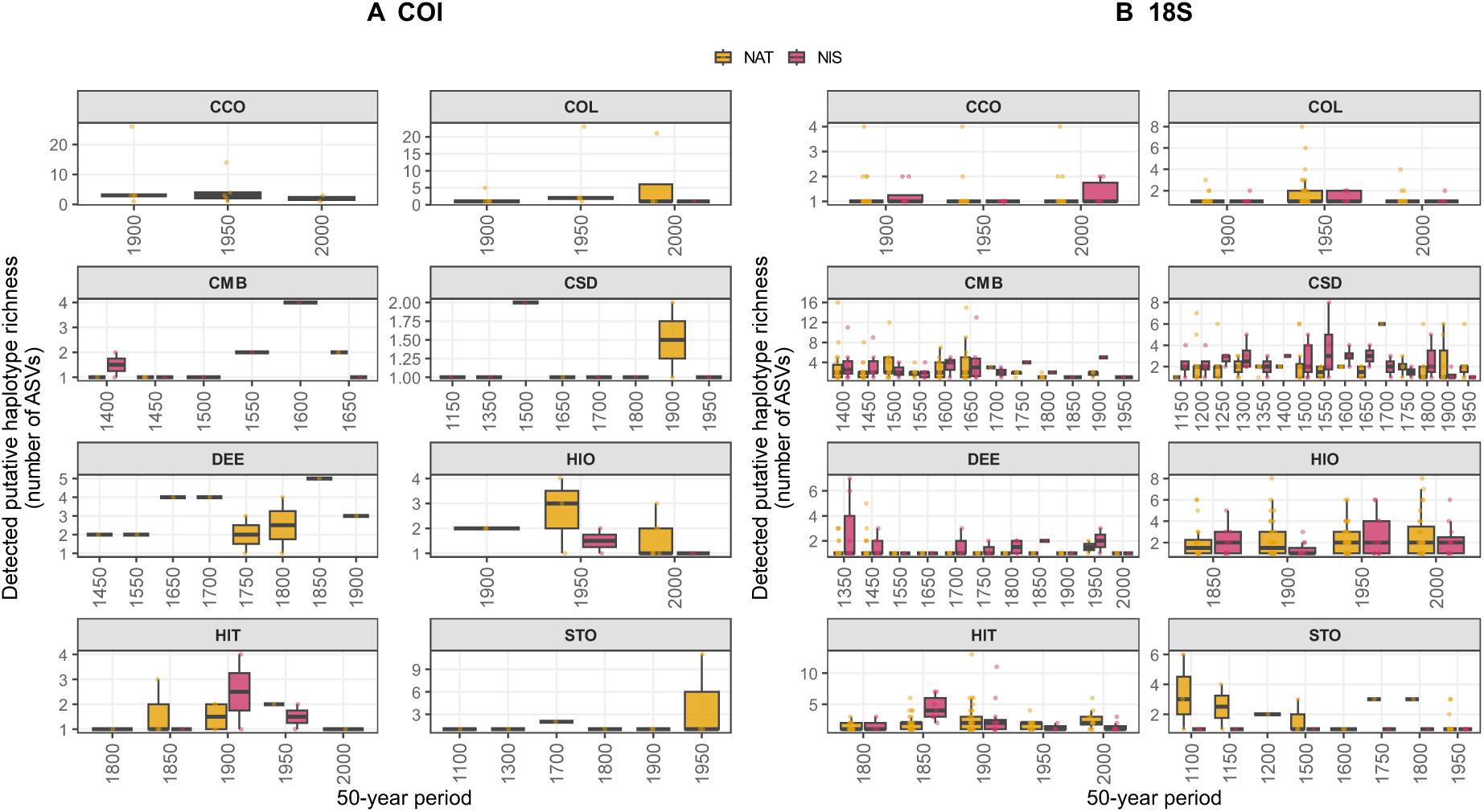
Temporal variation in detected putative haplotype richness of native and non-indigenous taxa across coastal sedimentary DNA archives. Detected putative haplotype richness, expressed as the number of ASVs detected per species within each 50-year period, is shown for (A) COI and (B) 18S across the eight study sites. Points represent individual species-period observations and boxplots summarize distributions for status groups represented by at least three species within a site and period. Native (NAT) and non-indigenous (NIS) taxa are shown separately. Statistical comparisons between NAT and NIS were performed within the same site and 50-year period using two-sided Wilcoxon rank-sum tests when both groups contained at least three species, with Benjamini–Hochberg correction across comparisons.

